# Reciprocal EGFR signaling in the Anchor Cell ensures precise inter-organ connection during *C. elegans* vulval morphogenesis

**DOI:** 10.1101/2021.06.16.448295

**Authors:** Silvan Spiri, Simon Berger, Louisa Mereu, Andrew DeMello, Alex Hajnal

## Abstract

During *C. elegans* vulval development, the uterine anchor cell (AC) first secretes an epidermal growth factor (EGF) to specify the vulval cell fates and then invades into the underlying vulval epithelium. Thereby, the AC establishes direct contact with the invaginating 1° vulF cells and attaches the developing uterus to the vulva. The signals involved and the exact sequence of events joining these two organs are not fully understood.

Using a conditional *let-23 egf receptor* (EGFR) allele along with novel microfluidic short- and long-term imaging methods, we discovered a specific function of the EGFR in the AC during vulval lumen morphogenesis. Tissue-specific inactivation of *let-23* in the AC resulted in imprecise alignment of the AC with the 1° vulval cells, delayed AC invasion and disorganized adherens junctions at the newly forming contact site between the AC and the dorsal vulF toroid. We propose that EGFR signaling, activated by a reciprocal EGF cue from the 1° vulval cells, positions the AC at the vulval midline, guides it during invasion and assembles a cytoskeletal scaffold organizing the adherens junctions that connect the developing uterus to the dorsal vulF toroid. EGFR signaling in the AC thus ensures the precise alignment of the two developing organs.

## Introduction

Signaling by epidermal growth factor receptor tyrosine kinases (EGFRs) controls a multitude of cellular processes during animal development (Segatto et al., 2011; Sisto et al., 2017; Yarden and Sliwkowski, 2001), with aberrant signaling resulting in various developmental defects and disease. Specifically over-activation of EGFR signaling, caused by mutations in the *egfr* gene itself or in genes encoding downstream signal transducers, is known to contribute to the formation and progression of various cancer types in humans (Hsu and Hung, 2016; Rajaram et al., 2017; Sibilia et al., 2007). However, the role of de-regulated EGFR signaling in cancer development is more complex than merely causing excess cell proliferation (Wee and Wang, 2017). Aberrant EGFR signaling in cancer cells promotes the formation of actin-rich protrusions, called invadopodia, and increases cell motility and invasion during metastatic growth, suggesting that secreted EGF ligands may act as chemo-attractants (Goswami et al., 2005; Yamaguchi et al., 2005). *egfr* knock-out mutations in mice, on the other hand, perturb the morphogenesis of various organs, resulting in defective branching morphogenesis and alveolarization of the lungs and defects in heart morphogenesis (Chen et al., 2016; Iwamoto and Mekada, 2006).

In contrast to mammalian genomes, which encode four EGFR family members and multiple types of epithelial growth factors (EGF), the *C. elegans* genome encodes a single EGFR tyrosine kinase, LET-23, and one EGF-like protein, LIN-3 (Aroian et al., 1990; Hill and Sternberg, 1992). As in higher organisms, LET-23 EGFR signaling in *C. elegans* controls many different developmental processes (Sundaram, 2006). In particular, EGFR signaling during the development of the egg-laying system, composed of vulva and uterus, has been studied in great detail (Gupta et al., 2012; Schindler and Sherwood, 2013). During vulval development, polarized LIN-3 EGF secretion from the AC towards the vulva precursor cells (VPCs) induces the 1° vulval cell fate by activating the EGFR/RAS/MAPK signaling pathway in P6.p, the nearest VPC, while at the same time acting as an attractive cue inducing VPC migration towards the AC (Grimbert et al., 2016). The VPCs adjacent to P6.p, P5.p and P7.p, receive less LIN-3 EGF signal and adopt the 2° vulval cell fate in response to a lateral Delta/Notch signal from P6.p, which inhibits the 1° fate (Berset et al., 2001; Greenwald et al., 1983; Yoo et al., 2004). The distal VPCs, P3.p, P4.p, and P8.p, only receive low doses of LIN-3 and Delta and adopt the 3° non-vulval cell fate. After the VPC fates have been specified, P5.p, P6.p and P7.p undergo three rounds of cell division, generating 22 vulval cells.

Although not strictly required for vulval induction, the VPCs and their descendants secrete a LIN-3 isoform that depends on processing by the ROM-1 Rhomboid protease to amplify the inductive AC signal via paracrine signaling (Dutt et al., 2004). In addition, LIN-3 expressed by the 1° descendants of P6.p specifies the uterine-vulval1 (uv1) cells later during the L4 stage (Chang et al., 1999; Saffer et al., 2011). During the L3 stage, after the vulval cell fates have been specified, the AC adopts an invasive phenotype and breaches the two basement membranes (BMs) separating the uterus from the developing vulva, establishing a direct connection between the two tissues (Sherwood and Sternberg, 2003). BM breaching requires polarization of the actin cytoskeleton within the AC towards the VPCs aligned on the ventral midline. AC polarity is induced by an UNC-6 Netrin signal from the ventral nerve cord (VNC), together with so far unidentified guidance cues from the 1° vulval cells (Sherwood and Sternberg, 2003; Ziel et al., 2008). Following invasion, vulval morphogenesis begins with the apical constriction and invagination of the 1° vulF cells. The other vulval cells migrate towards the vulval midline defined by the AC, extend circumferential protrusions and fuse with their contralateral partner cells, thereby forming a stack of seven syncytial rings, called the vulval toroids (Sharma-Kishore et al., 1999). Sequential contraction of the ventral and expansion of the dorsal toroids gives rise to a cylindrical lumen (Farooqui et al., 2012). Lastly, the vulF cells separate to expand the dorsal vulval lumen, and the AC occupies the space opened between the two vulF cells (Estes and Hanna-Rose, 2009). Consequently, the vulva is connected to the ventral uterus, and the AC fuses with the surrounding uterine cells to form the utse syncytium (Sapir et al., 2007). A robust connection between vulva and uterus is indispensable for egg-laying. However, the signals involved and the complete sequence of morphological changes that unite these two organs has not been fully resolved (Gupta et al., 2012).

While the function of the LET-23 EGFR pathway in specifying the vulval cell fates has been intensely investigated, possible roles of LET-23 signaling during vulval morphogenesis are not apparent, as they are likely masked by the earlier function of LET-23 during induction. The complete study of vulval development has additionally been hampered by the extended time period, over which the process occurs, and the often subtle phenotypic differences caused by aberrant vulval morphogenesis. We therefore used the microfluidic long-term imaging approach introduced by Berger *et al*. (Berger et al., 2021) to capture the entire process of vulval development in continuous time series. Furthermore, to quickly image large numbers of worms and therefore detect subtle phenotypic differences across different larval stages, we developed an adaptation of this imaging approach, which allows for short-term, high-throughput imaging. Combining these two methods, we were able to reveal for the first time the complete sequence of cell rearrangements orchestrated by the AC. Moreover, using a conditional *let-23* allele we show that LET-23 signaling in the AC is necessary to precisely position the AC and organize the formation of a new adherens junction between the AC and vulF toroid connecting the developing vulva to the uterus.

## Results

### LET-23 EGFR is expressed in in the Anchor Cell during vulval morphogenesis

To study the tissue-specific functions of EGFR signaling, we previously constructed a conditional, GFP-tagged *let-23 egfr* allele (*FRT::let-23::FRT::gfp(zh131)*) by sequential CRISPR/Cas9-mediated insertion of two Flippase (FLP) recognition target sites (FRT) flanking the tyrosine kinase domain of *let-23* along with a *gfp* reporter near the C-terminus (Konietzka et al., 2020). FLP-induced excision of the sequence flanked by the two FRT sites creates a *let-23* loss-of-function (*lf*) allele and simultaneously leads to the loss of the GFP signal.

While analyzing the expression pattern of the endogenous *let-23(zh131)* reporter, we not only observed the previously reported LET-23::GFP expression in the VPCs (Haag et al., 2014) and the uv1 cells (Chang et al., 1999), but also detected LET-23::GFP expression in the AC, from the onset of vulval invagination at the L4.0 stage (Mok et al., 2015) until the AC fused with the ventral uterine (VU) cells (**Fig. 1A-B’’**). The LET-23::GFP signal was diffusely localized in the AC near the contact site between the VPCs and uterus. To confirm the expression of LET-23::GFP in the AC, the *let-23(zh131)* allele was combined with a transgene expressing FLP driven by the AC-specific *lin-3* enhancer element and the minimal *pes-10* promoter (*zhEx614[P_ACEL-pes-10>_2xNLS-FLP-D5]*) (Hwang, 2004), hereafter referred to as *let-23AcKOEx*. In most *let-23AcKOEx* L4 larvae, the LET-23::GFP signal was absent in the AC without any obvious reduction in the VPCs (**Fig. 1C-C’’**), confirming expression in the AC in addition to the invaginating vulval cells during early vulval morphogenesis.

**Figure 1.**
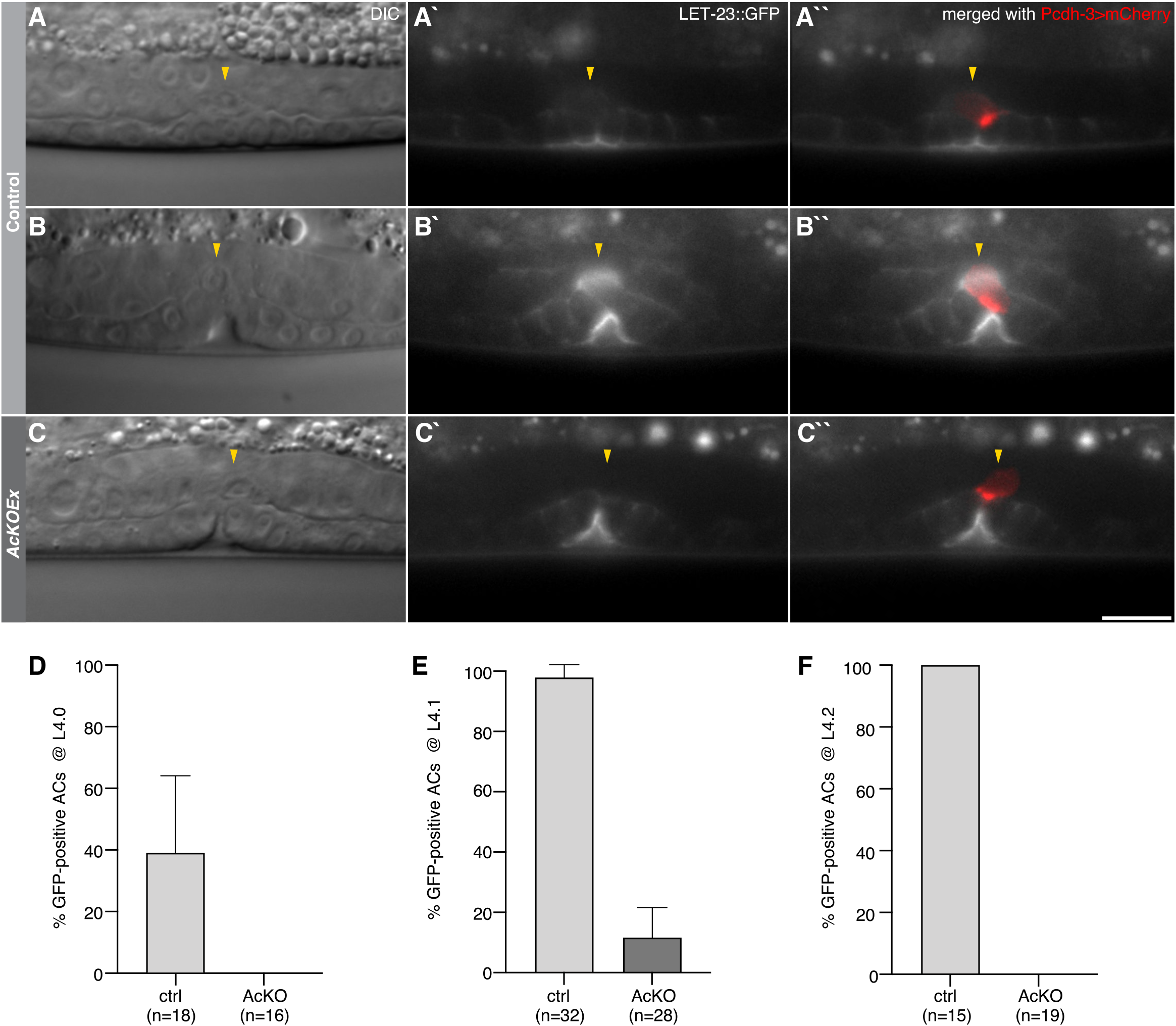
Endogenous LET-23::GFP expression in the AC. *let-23::gfp(zh131)* reporter expression at the (**A**) L4.0 and (**B**) L4.1 sub-stages of vulval development. (**C**) FLP recombinases-mediated deletion of *let-23::gfp(zh131)* using the *zhEx614[P_ACEL-pes-10>_2xNLS-FLP-D5]* extrachromosomal array, referred to as *let-23AcKOEx*. (**A-C**) DIC images of the mid-sagittal planes, (**A’-C’**) LET-23::GFP signal in gray and (**A’’-C’’**) LET-23::GFP signal overlaid with the *qyIs50[P_cdh-3>_mCherry::moeABD]* AC marker in red. Yellow arrowheads indicate the position of the AC. See **suppl. Fig. S2** for additional examples and stages with the *zhIs146[P_ACEL-pes-10>_2xNLS-FLP-D5]* transgene. (**D-F**) Fraction of animals expressing LET-23::GFP in the AC without (ctrl) and with (AcKO) the *zhIs146[P_ACEL-pes-10>_2xNLS-FLP-D5]* transgene at the (**D**) L4.0, (**E**) L4.1 and (**F**) L4.2 sub-stages. Error bars indicate the 95% CI and numbers in brackets the animals scored per condition. Scale bar is 10 *μ*m.

Initial analysis of vulval morphogenesis in *let-23AcKOEx* animals revealed a number of subtle and transient morphological defects, highlighting the importance of studying the dynamics of this process in real time as well as the necessity to rapidly image large numbers of worms.

### Short- and long-term microfluidic *C. elegans* imaging

Long-term *C. elegans* imaging was accomplished as described by Berger *et al*. (Berger et al., 2021), continuously imaging worms from the early L3 stage up to adulthood, capturing the entire process of vulval development. Briefly, worms were housed in a parallel array of channels, continually fed and actively immobilized during image acquisition using a hydraulic valve. Channel dimensions are designed to confine animals only in height to maintain a stable orientation and animal identity, but otherwise remaining free to grow and molt. Development on-chip, unlike on conventional agar-pads, occurs reliably and with minimal negative effects on the observed process (Berger et al., 2021).

The acquisition of z-stacks of selected worms at single time points, here termed short-term imaging, is traditionally accomplished by immobilizing animals through compression on agar-pads and by adding tranquilizing agents. While simple in construction, agar pad-based imaging is typically laborious with significant time spent on localizing and imaging randomly oriented animals. In the past, several microfluidic strategies for imaging of large worm populations have been introduce as an alternative. For example, Chung *et al*. introduced a sequential imaging method, where worms are automatically loaded into a single device (Chung *et al*. 2008, San-Miguel *et al*. 2016), while Mondal *et al*. (Mondal *et al*. 2016) proposed a parallel screening approach where several thousand animals are trapped in a single large device and subsequently imaged.

Both these methods allow imaging of large worm populations, but require dedicated microscope setups and peripherals (e.g. pumps, solenoid valves or dedicated control electronics), rendering adaptation of either method as a routine imaging tool difficult. Here, we developed a new parallel short-term imaging method, which is quick and easy to use for routine imaging, compatible with any microscope setup, without need for additional equipment and therefore suitable as an effective replacement for agar-pad immobilization in routine imaging. Compared to the original long-term imaging device (Berger et al., 2021), the short-term device layout is simplified with an array of 49 parallel trap channels between a single inlet and outlet (**Fig. 2A**). Immobilization in these devices is achieved passively through a combination of channel height relative to width (ratio H/W 0.7-0.8), with channels closely following the size and shape of a trapped worm at the desired stage, leaving little to no room for animal motion or growth. No active immobilization is employed, but immobilization can be further improved through addition of tetramisole or other chemical tranquilizing agents. Device dimensions for late L3 stage and mid-late L4 stage animals were chosen as 500×20×25*μ*m (type L3s) and 650×22×30*μ*m (type L4s) (LxHxW) respectively. Channel cross-section is kept constant, only tapering at the head and tail region, stabilizing these parts of the worm body and preventing worms from leaving the channel once loaded. All worms are trapped at the end of each channel, in a well-defined imaging region, by a drastic change in height (**Fig. 2A-B**, red). Thanks to the regular arrangement of the trap array and the straight worm orientation, images of multiple animals can be acquired in parallel at high-throughput with little time spent on localizing animals prior to image acquisition. Channel spacing was chosen such that in the field of view (FOV) of a sCMOS camera with an 18.7mm diagonal chip area, at 100X magnification two, at 60X three and at 40X five animals can be simultaneously imaged, with all 49 animals imaged in as few as 10 FOVs (**Fig. 2D)**. Acquisition speed may further be increased with eight identical devices fabricated on a single microscope slide, allowing imaging of up to 392 animals in quick succession, limited only by the microscopes acquisition speed (**Fig. 2 E**).

**Figure 2.**
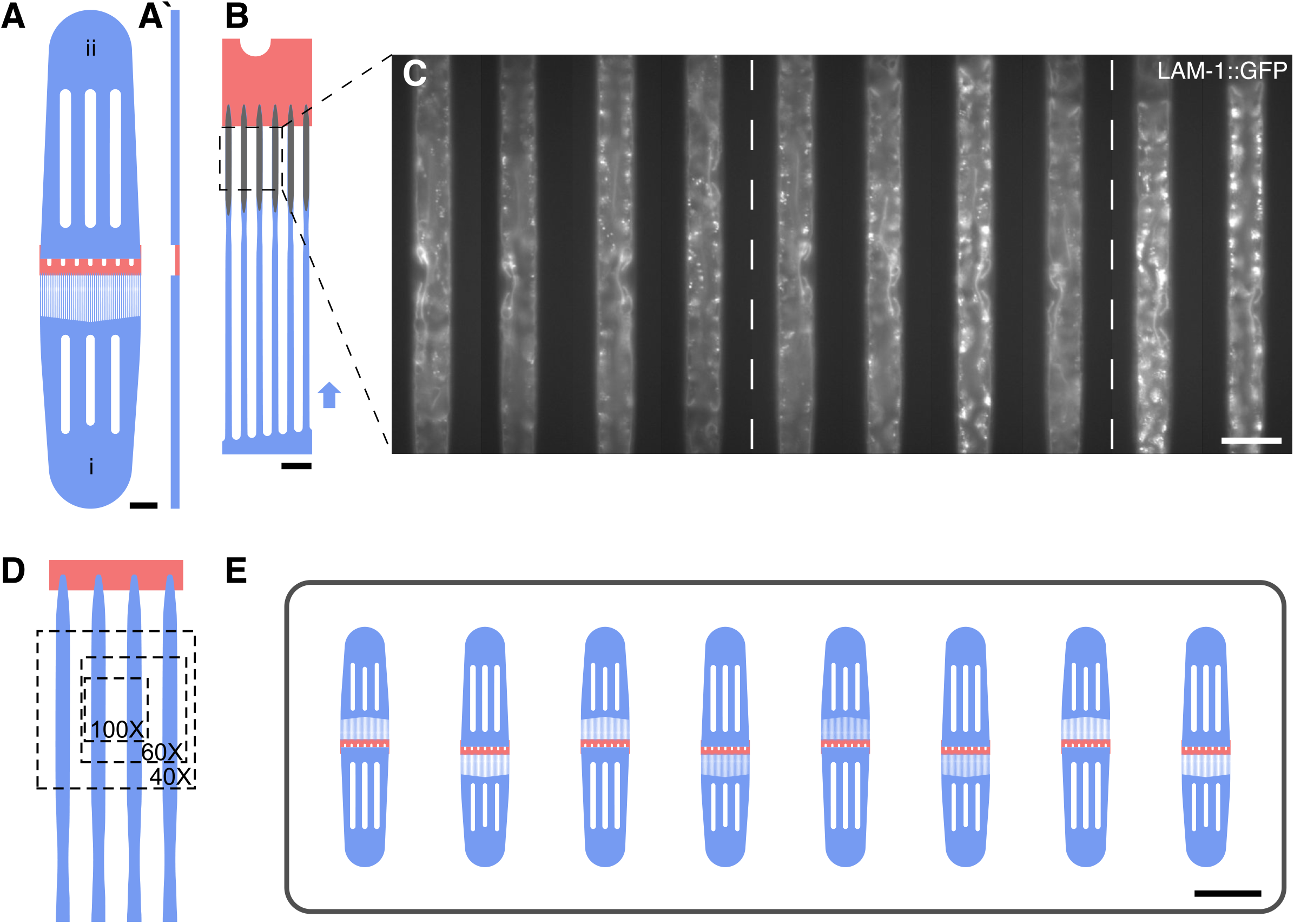
Microfluidic short-term imaging device. (**A**) Schematic overview of a single short-term imaging device. Worms are loaded from (i) with the excess fluid drained via the outlet (ii). (**A’**) Cross-sectional view (not to scale). The trap region (blue, bottom) is separated from the outlet region (blue, top) via a low height region (red) effectively preventing worms from passing from one device compartment into another. (**B**) Magnified device view. Worms are manually loaded in direction of the arrow, such that each of the 49 trap channels is filled with a single worm. Worm immobilization and orientation is achieved through carefully tuned device dimensions relative to the developmental stage studied, with the low height region (red) stopping worms at the end of the trap channel. Once loaded, worms are kept in the imaging by the tapered section at the tail end of each animal. (**C**) Representative images acquired on-chip showing a basement membrane marker *qyIs127[P_lam-1>_lam-1::gfp]* at the early to mid-L4 stage. All worms are confined at the end of the trap channel and lined up in parallel, such that multiple worms may be imaged in one field of view (4-5 animals at 40X magnification, dashed box) and multiple fields of view may be imaged by moving along the device. (**D**) Magnified view of the parallel imaging scheme. Overlay shows the ROI achievable with different magnifications. (**E**) Schematic of a single microscope slide with 8 individual devices capable of housing 392 animals at the same time. Scale bars are (**A**) 1000*μ*m, (**B**) 100*μ*m, (**C**) 50*μ*m and (**E**) 5000*μ*m.

Device operation is even simpler than in the long-term imaging variant, such that a large number of worms can be loaded in only a few minutes. Synchronized worms are washed off the NGM plates, loaded using a syringe connected to the inlet of a specific device through a short piece of tubing (see **Extended methods** for a detailed step-by-step protocol). Worms passively fill the trap channels one-by-one, as device dimensions are chosen close to the desired worm stage, resulting in a loading efficiency of approximately 95% with all loaded animals trapped in lateral orientation. Once loaded, the tubing is removed and the device can easily be mounted on any microscope (upright and inverse). Importantly, animals remain viable for several hours on-chip without adverse effects of the device on worm health or morphology. In fact, short-term imaging devices may even be used for acquisition of 1 to 2 hour-long timelapse sequences, with trapped worms feeding on bacteria loaded along with the animals.

All devices are fabricated from highly transparent PDMS and mounted on cover glass, such that images may be acquired using high-magnification and high NA objectives, and in combination with most commonly used imaging modalities (widefield and confocal microscopy) without any noticeable drop in image quality compared to animals imaged on agar pads (**Fig 2C**). Consequently, all images shown in this study, except for those in **Fig. 1** and **S1**, were acquired using either the short- or long-term microfluidic imaging devices.

### LET-23 expression in the AC ensures precise alignment of the AC with the invaginating 1° vulval cells

LIN-3 EGF is secreted by the 1° vulval cells as a relay signal to ensure robust vulval induction during the L3 stage (Dutt et al., 2004), and to specify the uterine uv1 cell fate later during the L4 stage (Chang et al., 1999). We therefore investigated if LET-23 EGFR might also function in the AC receiving a reciprocal LIN-3 signal from the 1° VPCs.

For the following experiments, a stable *let-23AcKO* mutant strain was generated by combining the conditional *let-23(zh131)* allele with a single-copy integrated FLP transgene *zhIs146[P_ACEL-pes-10>_2xNLS-FLP-D5]*, hereafter referred to as *let-23AcKO* (Frøkjær-Jensen et al., 2008). To probe the activity and specificity of the FLP expression, the *zhIs146* transgene was crossed with the heat shock-inducible FLP activity reporter *bqSi294[P_hsp16.41>_FRT::mCherry::his-58::FRT::GFP::his-58]* (Muñoz-Jiménez et al., 2017), which in cells with an active FLP, converts mCherry fluorescence into a nuclear GFP signal upon heat-shock. The *zhIs146* transgene induced conversion of the *bqSi294* reporter in the AC with 100% (n=20) efficiency (**Fig. S1A,C**). In 80% (n=20) of the animals, one or two of the adjacent VU descendants were also converted, probably due to weak activity of the *ACEL-pes-10* enhancer/promoter during AC/VU specification at the early L2 stage (**Fig. S1A,D**). By contrast, FLP activity of the *zhIs146* transgene was never observed in the VPCs (**Fig. S1A,B**).

Faint LET-23::GFP expression in the AC was observed beginning at the L4.0 substage (Mok et al., 2015) in 39% (n=18) of control animals lacking the *zhIs146* transgene, compared to 0% (n=16) of *let-23AcKO* mutants (**Fig. 1A,D** and **Fig. S2A-B’**). During the following L4.1 sub-stage, 98% (n=32) of the control animals compared to 12% (n=28) of *let-23AcKO* mutants showed AC expression of LET-23::GFP (**Fig. 1B,E** and **Fig. S2C-D’**), and in L4.2 larvae, the last sub-stage before the AC fuses with the utse, AC expression was observed in all controls (n=15) and in none of the *let-23AcKO* mutants (n=19) (**Fig. 1F** and **Fig. S2E-F’**). No LET-23::GFP expression in other uterine cells besides the AC could be detected before the L4.3 stage, when LET-23::GFP expression was first observed in the uv1 descendants of the VU cells in 76% (n=13) of control animals (**Fig. S2I**). uv1 expression of LET-23::GFP was absent in 80% (n=42) of *let-23AcKO* animals by the L4.4 stage, probably due to earlier activity of *zhIs146* in the VU cells, as mentioned above (**Fig. S2G,H,J** and **Fig. S1A,D**). Since the AC is the only cell in the ventral uterus that expresses LET-23 during the period of vulval invagination and toroid formation (before L4.3), the phenotypes induced by the ACEL-FLP transgene described in the following are most likely due to the loss of LET-23 function in the AC.

We analyzed vulval morphogenesis in *let-23AcKO* mutants using the endogenous cadherin reporter *hmr-1::gfp(cp21)* to visualize the adherens junctions together with the actin marker *qyIs50[P_cdh-3>_mCherry::moeABD]* labeling the AC (Marston et al., 2016; Ziel et al., 2008). This analysis revealed a displacement of the AC from the vulval midline in *let-23AcKO* mutants at the L4.0 sub-stage, the first time point in which AC expression of LET-23::GFP was detected (**Fig. 3A,B**). We quantified AC positioning in each animal by calculating an alignment ratio (R_A_), defined as the relative position of the AC mid-point to the invaginating vulval cells (with R_A_ = 0.5 indicating perfect centering), as well as by measuring the absolute distance (Δ) between the AC and vulF mid-points (**Fig. 3C** and **Materials and Methods**). In control animals lacking the *zhIs146* transgene, the mean R_A_ was 0.462±0.028 (n=113), indicating close alignment of the AC with the invaginating vulval cells. By contrast, AC alignment in *let-23AcKO* mutants was less precise with a mean R_A_ value of 0.435±0.04 (n=134) and a higher variability of R_A_ (**Fig. 3D**). Also, the absolute distance Δ showed an increased variance in *let-23AcKO* mutants (**Fig. 3E**, s.d. control: ±1.584 *μ*m; s.d. *let-23AcKO*: ±2.380 *μ*m).

**Figure 3.**
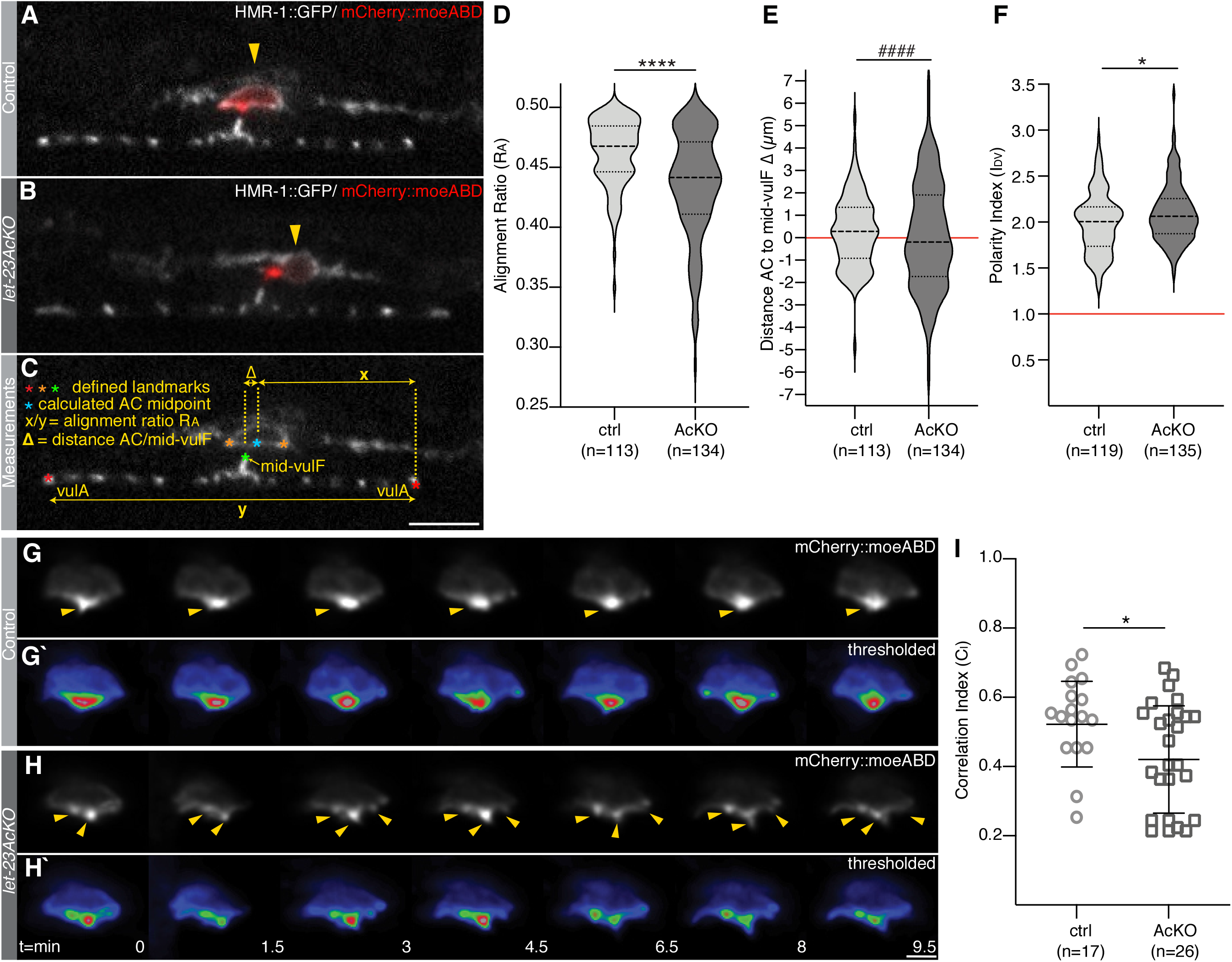
Imprecise AC alignment and disorganized actin protrusions after AC inactivation of LET-23. (**A**) Mid-sagittal planes of HMR-1::GFP (*cp21*) adherens junction marker in white overlaid with the *qyIs50[P_cdh-3>_mCherry::moeABD]* AC marker in red in a *let-23(zh131)* control animal lacking the *zhIs146[P_ACEL-pes-10>_2xNLS-FLP-D5]* transgene and (**B**) a *let-23AcKO* animal at the L4.0 stage. (**C**) Illustration of the landmarks (red, orange and green asterisks) selected when measuring the AC alignment index R_A_ (ratio of the shorter x to y) and the absolute AC to vulF distance Δ. The blue asterisk indicates the AC center calculated as the mid-point between the two orange asterisks. (**D**) Violin plots of the AC alignment index R_A_ in control (ctrl) and *let-23AcKO* animals at the L4.0 stage. (**E**) Violin plots of the absolute distance AC to mid-vulF in control and *let-23AcKO* animals. (**F**) Violin plots of the dorso-ventral AC polarity index I_DV_ measured with the *qyIs50[P_cdh-3>_mCherry::moeABD]* AC marker as described (Mereu et al., 2020). (**G)** Time series of the *qyIs50[P_cdh-3>_mCherry::moeABD]* actin reporter in the AC in a control and (**H**) a *let-23AcKO* larva at the L4.0 stage acquired at 30-second intervals, every third frame is shown (see **Movies S1,2** for all time points). Yellow arrowheads in (**G**) point to the single AC protrusion formed at the mid-line in a control and in (**H**) to the multiple AC protrusions formed in *let-23AcKO* animals. (**G’**) and (**H’**) show heat-maps of the thresholded AC images used to calculate the frame-to-frame correlation indices C_I_. (**I**) Average correlation indices C_I_ measured in the AC of control and *let-23AcKO* L4.0 animals. Bars in (**I**) indicate the mean values and standard deviation. Dashed lines in the violin plots in (**D-F**) indicate the median values and the dotted lines the upper and lower quartiles. Statistical significance was calculated in (**D**) and (**F**) with a t-test for independent samples of unequal variance and in (**I**) with a nonparametric, unpaired Man-Whitney test since the data were not normally distributed (* for p <0.05 and **** for p <0.0001). #### in (**E**) indicates the result of an F-test for variance, indicating unequal variance with p <0.0001. Numbers in brackets below each graph refer to the numbers of animals scored per genotype. Scale bars are 10 *μ*m in (**A**) and 5 *μ*m in (**H’**).

To test if the observed AC positioning defect was caused by a depolarization of the AC upon inactivation of LET-23, we measured the dorso-ventral polarity index (I_DV_) using the *qyIs50[P_cdh-3>_mCherry::moeABD]* reporter as described (Mereu et al., 2020 and **Material and Methods**). This analysis revealed a slightly increased, rather than decreased AC polarity index in *let-23AcKO* mutants, suggesting that LET-23 signaling is not necessary to polarize the AC (**Fig. 3F**).

Next, we performed rapid time-lapse imaging experiments to capture the dynamics of the actin cytoskeleton. For this purpose, we imaged the *qyIs50[P_cdh-3>_mCherry::moeABD]* actin reporter at 30 second intervals for a total of 10 minutes in animals trapped in the short-term device. In most control animals, the actin reporter signal was concentrated in a single dot on the ventral AC surface, directed towards the mid-point of the invaginating vulF cells. In the AC of *let-23AcKO* mutants, we observed several actin foci in multiple protrusions (**Fig. 3G-H’** and **Movies S1,2**). The dynamic changes of the AC were quantified through correlation analysis of consecutive time points, effectively assessing the degree by which AC morphology changes over time (**Material and Methods**). This analysis indicated a more dynamic rearrangement of the AC (i.e. lower correlation index C_I_) in *let-23AcKO* mutants (**Fig. 3I**).

In summary, our data indicate that LET-23 expression in the AC is required for the precise alignment of the AC with the invaginating vulval cells. LET-23 signaling might act by organizing the actin cytoskeleton in the AC protrusions directed towards the vulval midline.

### LET-23 signaling in the AC promotes basement membrane breaching

Over the course of vulval organogenesis, the AC breaches two BMs to establish a physical connection between the developing vulva and uterus (Sherwood and Sternberg, 2003). During this process, the AC is polarized and guided by an UNC-6 Netrin signal from the VNC toward the underlying vulval tissue (Ziel et al., 2008). In wild-type animals, the BMs have been completely breached at the beginning of vulval invagination at the L4.0 sub-stage, as AC invasion normally occurs during mid-L3 (the Pn.pxx stage). In *unc-6(lf)* mutants, the BMs are only breached in around 30% of the animals at the L4.0 stage, but by the L4.4 sub-stage BMs breaching has occurred in 80% of the animals, indicating delayed AC invasion (Ziel et al., 2008). It has therefore been proposed that UNC-6 Netrin acts in parallel with one or several additional diffusible guidance cues secreted by the 1° vulval cells (Sherwood and Sternberg, 2003).

As loss of LET-23 function affects AC alignment and actin dynamics, we tested if LET-23 signaling may play a role in guiding the invading AC. For this purpose, we observed BM breaching in *let-23AcKO* mutants carrying an *mCherry*-tagged *lam-1 laminin* reporter (*qyIs127[P_lam-1>_lam-1::mCherry]*) labelling the BMs (Ihara et al., 2011). *let-23AcKO* single mutants exhibited a slight but significant delay in BM breaching at the Pn.pxx stage, as only 64% of *let-23AcKO* animals (n=90) showed a gap in their BMs, compared to 83% of control animals (n=121) (**Fig. 4A,B,G**). However, by the L4.0 substage the BMs were breached in all *let-23AcKO* mutants (n=107) (**Fig. 4G**). This suggested that, while not being essential for invasion, LET-23 signaling in the AC might cooperate with other guidance cues to promote BM breaching. We thus analyzed AC invasion in *let-23AcKO; unc-6(ev400lf)* double mutants. Only 11% (n=89) of *let-23AcKO; unc-6(ev400lf)* double mutants at the L4.0 sub-stage exhibited BM breaching, compared to 29% (n=59) of *unc-6(ev400lf)* single mutants (**Fig. 4 C,D,H**). This effect persisted at later stages, with 40% (n=63) of *let-23AcKO; unc-6(ev400lf)* double mutants showing BM breaching by the L4.4 stage, compared to 68% (n=25) of *unc-6(ev400lf)* single mutants (**Fig. 4E,F,H**).

**Figure 4.**
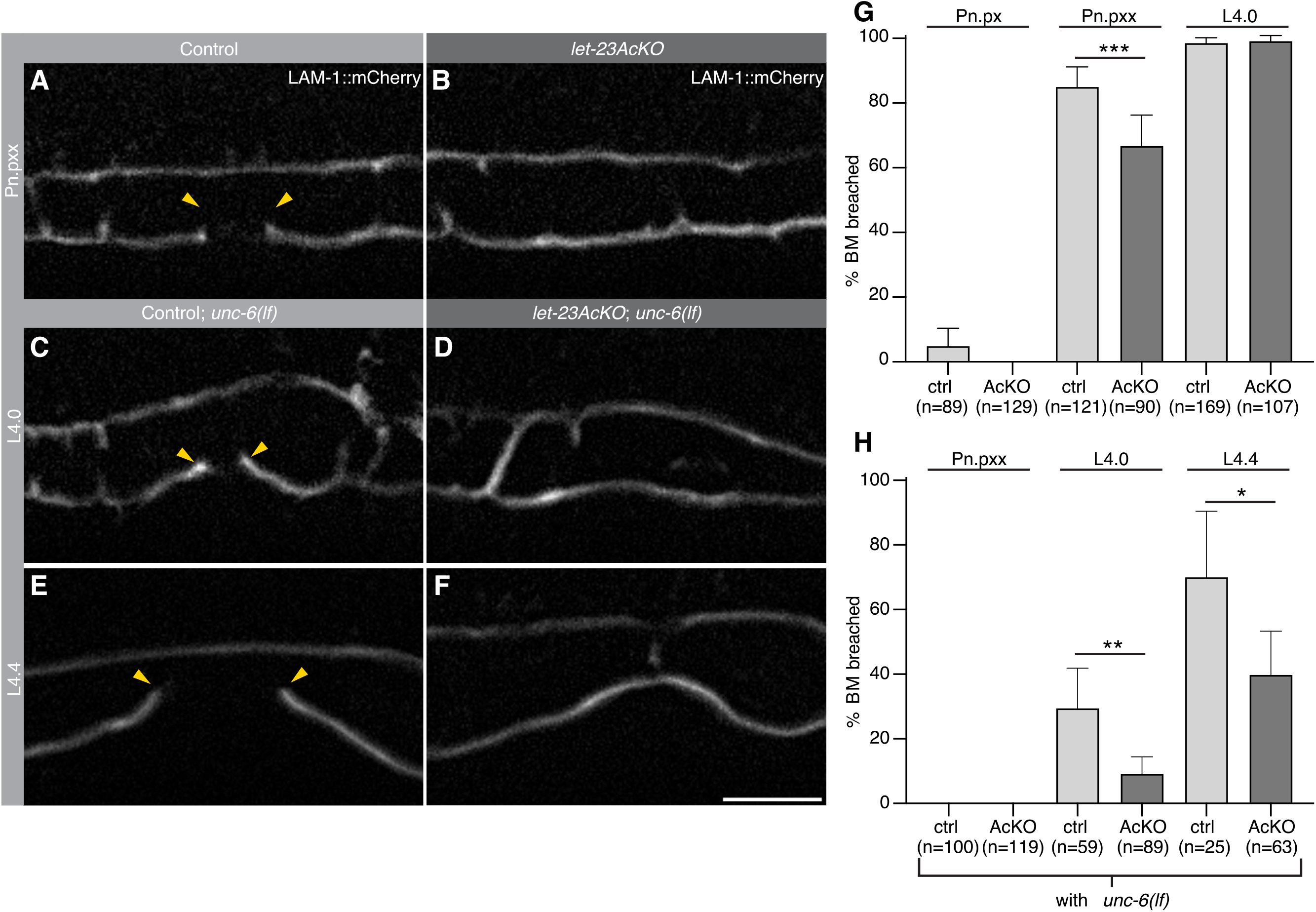
Basement membrane breaching defects after AC specific LET-23 inactivation. (**A**) Mid sagittal planes showing expression of the *qyIs127[P_lam-1>_lam-1::mCherry]* BM marker in a *let-23(zh131)* control animal lacking the *zhIs146[P_ACEL-pes-10>_2xNLS-FLP-D5]* transgene and (**B**) a *let-23AcKO* animal at the Pn.pxx stage of L3 stage. (**C**) LAM-1::mCherry expression in *let-23(zh131); unc-6(ev400lf)* and (**D**) *let-23AcKO; unc-6(ev400lf)* animals at the L4.0 stage (beginning of invagination) and (**E, F**) at the L4.4 stage (completed invagination). The yellow arrowheads in (**A, C** & **E**) indicate the extent of BM breaching. (**G**) Frequency of BM breaching at the indicated stages in control (ctrl) and *let-23AcKO* animals. (**H**) Frequency of BM breaching at the indicated stages in control; *unc-6(ev400lf)* and *let-23AcKO; unc-6(ev400lf)* animals. Error bars indicate the 95% CI and numbers in brackets the animals scored per condition. Statistical significance was calculated using a non-parametric, unpaired Mann-Whitney test (* for p <0.05, ** p <0.01 and *** p <0.001). Scale bar is 10 *μ*m.

We conclude that LET-23 signaling in the AC transduces a guidance cue that cooperates with the UNC-6 Netrin signal secreted from the VNC to achieve efficient BM breaching. Since LIN-3 EGF is only expressed by the 1° vulF cells at this stage (Chang et al., 1999; Saffer et al., 2011), it seems likely that LET-23 in the AC receives the LIN-3 signal from the invaginating vulval cells.

### Inactivation of LET-23 in the AC causes disorganized vulval-uterine junctions

AC invasion and alignment are crucial to initiate dorsal lumen formation during the subsequent stages of vulval morphogenesis (Estes and Hanna-Rose, 2009). After invasion, the AC remains positioned at the vulval midline in a pocket surrounded by the invaginating vulF cells. If the AC fails to invade into the vulval tissue or if it is displaced, an abnormal dorsal vulval lumen will form (Estes and Hanna-Rose, 2009). We therefore investigated the consequences of the early AC positioning defect observed in *let-23AcKO* mutants at the L4.0 stage on vulval lumen morphogenesis. To obtain detailed insights into the dynamics of vulval morphogenesis, we used long-term imaging devices (type L2-A, Berger *et al*. 2021), which allowed seamless imaging of the same animal from the early L3 stage until adulthood. Vulval morphogenesis was observed in control and *let-23AcKOEx* animals carrying the *hmr-1::gfp(cp21)* and *qyIs50[P_cdh-3>_mCherry::moeABD]* reporters, outlining the adherens junctions and AC, respectively (**Movies S3,4**).

These experiments showed that the AC in control animals remained fixed after it had made initial contact with the invaginating vulF cells (**Fig. 5A**). By contrast, the AC position in *let-23AcKOEx* mutants was highly variable, causing the connected toroids to rock back and forth along the anterior-posterior axis (**Fig. 5B**). In the following, we assed robustness of vulval morphogenesis by measuring vulA migration and found that the distance between the outer vulA junctions decreased at a similar rate in both strains, indicating that the migration of the vulval cells towards the midline is not affected by loss of LET-23 function in the AC (**Fig. 5C,D**). The distance between the AC and vulF midpoints (Δ) relative to vulA distance, used as an indicator of developmental time, remained large and variable in *let-23AcKOEx* mutants (n=12), while it continually decreased in control animals (n=9) prior to AC fusion (**Fig. 5E,F** and **Fig. S3A,B**).

**Figure 5.**
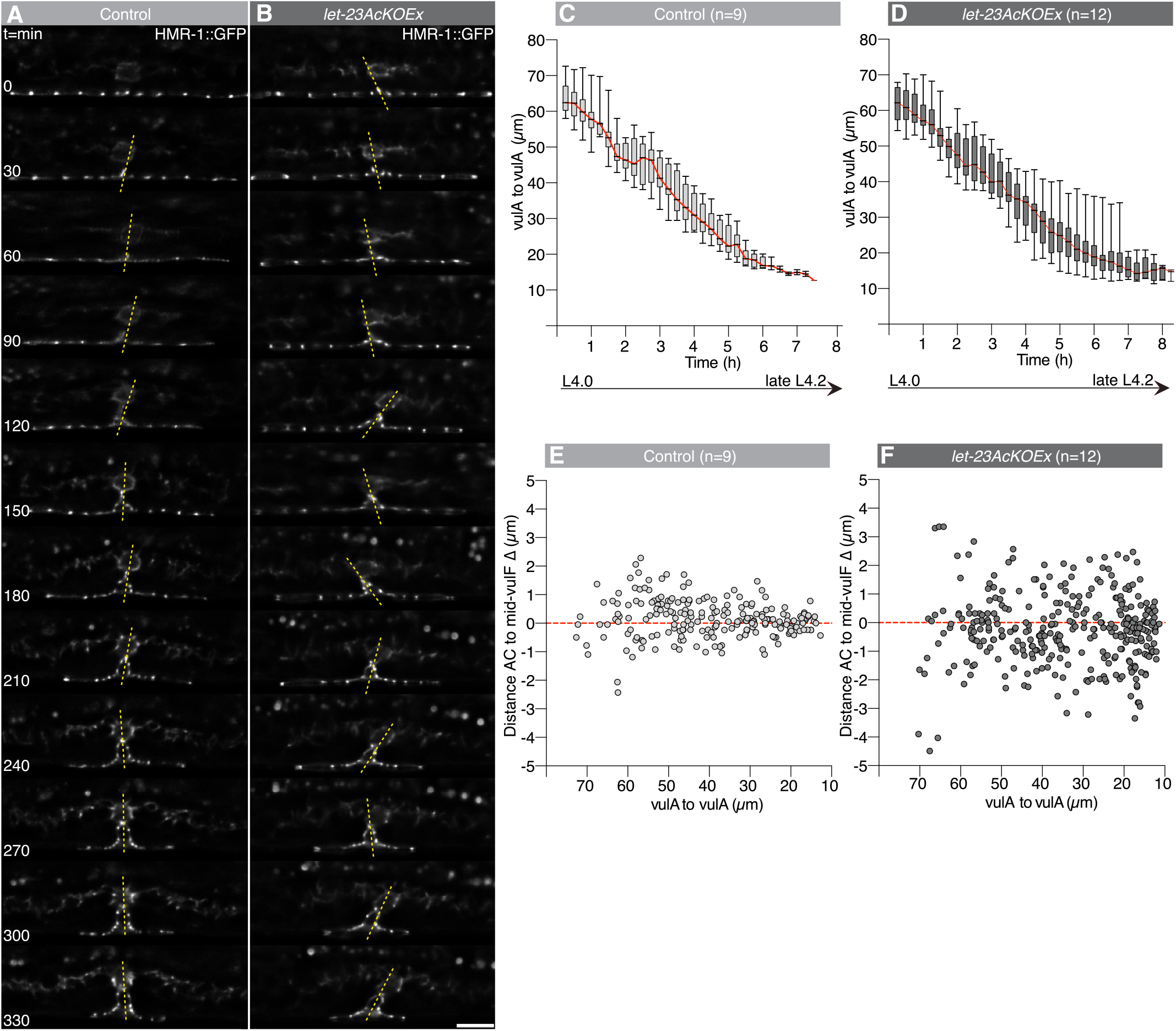
Variable AC position during vulval invagination after LET-23 inactivation. (**A**) Time series showing the mid sagittal plane for the HMR-1::GFP (*cp21*) adherens junction marker between the L4.0 and late L4.2 stages in a *let-23(zh131)* control and (**B**) *let-23AcKOEx* animal. Yellow dashed lines indicate the AC position relative to the vulval midline. Images shown at 30-minute intervals. (**C**, **D**) Box plots showing the changes in the distance between the outer vulA junctions over time, used as a measure for the migration of the vulval cells to the midline. The red lines connect the medians throughout the timepoints. (**E**, **F**) Dot plots showing the absolute AC to mid-vulF distances (Δ) relative to the vulA to vulA distance. Error bars show minimum and maximum values, boxes the 25% to 75% quartiles and black horizontal lines the median values. The numbers in brackets indicate the animals scored per genotype. See also **suppl. Fig. S3** for the individual measurements over time for each animal. Scale bar is 10 *μ*m.

After generating 3D reconstructions of the time-lapse recordings, we noticed that bright HMR-1::GFP dots appeared in the AC body dorsal to the actin-rich AC protrusion, which first contacts the vulF cells, indicating that new adherens junctions were assembling in the AC (**Fig. 6A-A’’**, yellow arrowheads frames 1-2, also **suppl. Fig S4A**). Subsequently, a ring-shaped structure emerged from these adherens junctions (**Fig. 6A-A’’**, yellow arrowheads frames 3-4), which contracted as it joined with the adherens junctions on the dorsal surface of the vulF toroid (**Fig. 6A-A’’**, yellow arrowheads frames 5-6). At the same time, the AC protrusion retracted and the AC began to fuse with the utse, as indicated by diffusion of the *mCherry::moeABD* signal into adjacent uterine cells. The ring of adherens junctions subsequently expanded and the dorsal vulval lumen was opened in all control animals (**Fig. 6A-A’’**, yellow arrowheads frames 7-10 and **Movie S5**). During these events, a distinct actin-rich domain remained localized at the interphase between vulF and the fusing AC in eight out of nine control animals (**Fig. 6A’**, yellow arrowheads frames 6-8).

**Figure 6.**
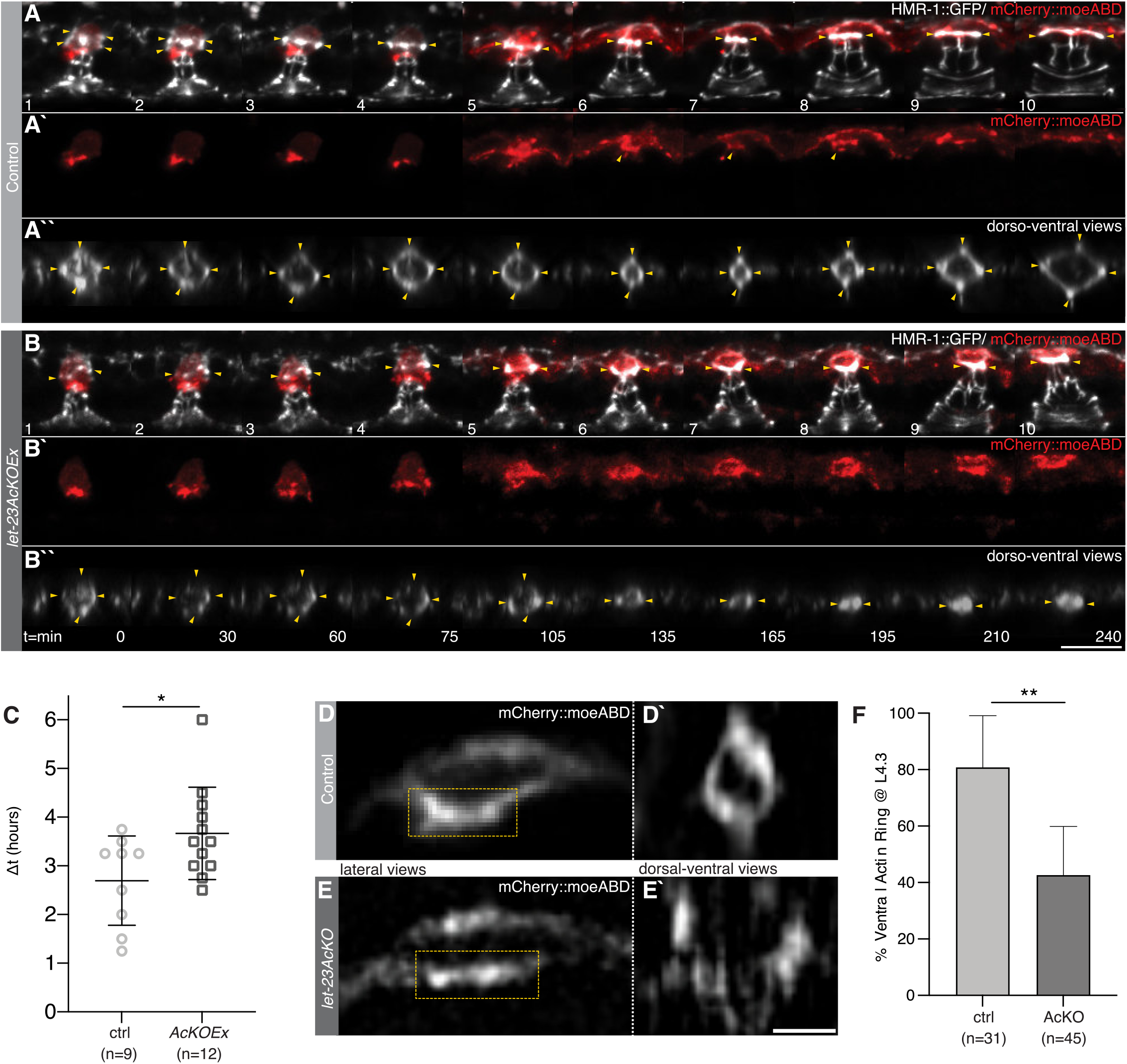
Vulval toroid morphogenesis and actin ring assembly after AC specific LET-23 inactivation. (**A-B**) Time series of the HMR-1::GFP (*cp21*) adherens junction marker in white overlaid with the *qyIs50[P_cdh-3>_mCherry::moeABD]* AC marker in red in a (**A**) *let-23(zh131)* control (ctrl) and (**B**) a *let-23AcKO* animal between the L4.1 and L4.4 stages (end of dorsal lumen expansion in ctrl). **A1-10** and **B1-10** show maximum intensity z-projections at the indicated time points. (**A’1-10** and **B’1-10**) show the P_cdh-3>_mCherry::moeABD actin marker in the AC. Yellow arrowheads in **A1-10** and **B1-10** point to newly forming adherens junctions in the AC and in **A’6-8** to the actin cytoskeleton near the adherens junctions. **A’’1-10** and **B’’1-10** show dorso-ventral views (cropped xz-projections) of the AC-vulF interphase, yellow arrowheads indicating the newly forming adherens junctions. (**C**) Time from AC fusion to the start of dorsal lumen expansion, between the L4.2 and L4.3 stages. Δt was defined as the time period between the last time point before the beginning of AC fusion (**A4**), indicated by the diffusion of the mCherry::moeABD signal into adjacent uterine cells, and the first time point when an expansion of the dorsal adherens junction ring was observed (**A’’8**). See **Movies S3,4** for all time points and **suppl. Fig. S4** for additional examples. (**D-E**) Confocal images of the P_cdh-3>_mCherry::moeABD actin marker in the AC of (**D**) control and (**E**) *let-23AcKO* animals at the L4.3 stage. (**D’**) and (**E’**) show dorso-ventral views (xz-projections) of the actin signal at the AC-vulF interphase in the regions highlighted in (**D**) and (**E**) by yellow dashed boxes. See **suppl. Fig. S5** for additional examples. (**F**) Fraction of control and *let-23AcKO* animals at the L4.3 substage containing a defined actin (semi)circle at the AC-vulF interphase. Error bars in (**C**) and (**F**) indicate the 95% CI. Statistical significance was calculated in (**C**) with a t-test for independent samples of unequal variance and in (**F**) with a non-parametric, unpaired Mann-Whitney test (* for p <0.05 and ** for p <0.01). Numbers in brackets below each graph refer to numbers of animals scored per genotype. Scale bars in (**B’’10**) 10 *μ*m and (**E’**) 2 *μ*m.

In *let-23AcKOEx* mutants, the AC protrusion did initially contact the vulF cells and bright HMR-1::GFP dots also appeared in the AC, but the adherens junctions were disorganized and no clearly defined ring structure could be seen in eight out of twelve animals (**Fig. 6B** and **Movie S6**, also **suppl. Fig S4B**). Despite the absence of a properly organized adherens junction ring in the AC, the expansion of the dorsal vulval lumen did occur in *let-23AcKOEx* mutants, but it appeared to be delayed. To quantify the speed of dorsal lumen expansion, we measured the time from the last frame just prior to AC fusion to the point, when the adherens junctions started to expand (Δt) (e.g. **Fig. 6A-A’’**, frames 4 and 8). The average Δt was 2.69±0.91 hours (n=9) in control and 3.66±0.94 hours (n=12) in *let-23AcKOEx* animals (**Fig. 6C)**.

In summary, time-lapse observation of vulval morphogenesis revealed that LET-23 signaling is necessary to assemble the newly forming adherens junctions in the AC into a ring-shaped structure, such that the AC and the vulF toroid can be properly joined before the dorsal vulval lumen expands.

### LET-23 signaling establishes a ring-shaped actin scaffold at the AC-vulF interphase

To further examine the connection between the AC and vulF toroid, we recorded confocal images of the F-actin reporter *qyIs50* at the stage shortly before the AC fuses to form the utse and generated xz-projections along the plane of the AC-vulF junction (e.g. **Fig. 6A’** frame 6). This analysis revealed in 81% (n=31) of control animals the existence of a distinct, actin-rich ring around the contact site between the ventral AC membrane and the dorsal surface of the vulF toroid (**Fig. 6D,D’,F** and **suppl. Fig. S4**). In 60% of *let-23AcKO* mutants (n=45), this actin-rich structure was either absent or only barely recognizable (**Fig. 6E-F** and **suppl. Fig. S5**).

Thus, LET-23 signaling is necessary to reorganize the AC cytoskeleton and assemble an actin-rich scaffold, which may serve as a template to organize the newly forming adherens junctions at the AC-vulF contact site.

## Discussion

### Microfluidic *C. elegans* handling

Both, the previously introduced long-term image devices (Berger et al., 2021) as well as the short-term imaging variant introduced here open a number of new possibilities. As described by Berger *et al*. (Berger et al., 2021) the long-term imaging devices allow the observation of various processes occurring over multiple developmental stages within the same animal, resulting in a better understanding of developmental dynamics. The short-term devices introduced here complement the possibility to study single animals over time with the ability to quickly image large numbers of worms at considerably higher throughput than possible using traditional agar pad immobilization without affecting worm viability or image quality. Devices for any developmental stage may be created following the simple design parameters outlined here, streamlining routine image acquisition as well as allowing the capture of subtle phenotypic variations only visible in large populations or at specific developmental stages. In the future, the short-term imaging devices may be combined with automated image acquisition tools for a variety of forward and reverse genetic screening experiments.

### LET-23 EGFR acts in the AC during early vulval morphogenesis

The Nematode *C. elegans* has served as a powerful *in vivo* model to dissect the various roles the EGFR signaling pathway plays during animal development and adulthood (Chamberlin and Sternberg, 1994; Clandinin et al., 1998; Konietzka et al., 2020; Pu et al., 2016; Van Buskirk and Sternberg, 2007).

In particular, EGFR signaling is essential during the development of the *C. elegans* vulva, the egg-laying organ of the hermaphrodite. Here, we provide evidence for a previously unknown role of EGFR signaling in the uterine AC during invasion and vulval morphogenesis. Using a conditional *let-23* knock-out allele, we found that inactivation of LET-23 EGFR in the AC and ventral uterine cells results in a defective interconnection between the vulval toroids and the ventral uterus. Even though the FLP driver line used is not specific to the AC, as it also induced recombination in the adjacent VU cells, the defects we have observed are most likely due to the loss of LET-23 EGFR activity in the AC, as it is the only cell in the ventral uterus expressing LET-23 during vulval invagination and toroid formation. Only at a later stage, LET-23 expression could be detected in the four uv1 cells that are specified during the mid-L4 stage by an EGF signal from the vulF cells (Chang et al., 1999). The uv1 cells join with the vulF toroid and utse only after the AC has connected with vulF and fused to form the utse, and animals lacking uv1 cells develop a normal vulval-uterine connection (Ihara et al., 2011; Johnson et al., 2009). Therefore, it is unlikely that the earlier defects in AC positioning and AC-vulF junction formation observed after inactivation of LET-23 are caused by a defect in uv1 fate specification or by undetectable LET-23 expression in other uterine cells. We can however not exclude the possibility that some of the defects observed later during dorsal lumen morphogenesis, such as the delay in dorsal lumen opening, may be caused by a lack of uv1 cells.

### Reciprocal EGFR signaling positions the AC and promotes BM breaching

The first phenotype we observed after inactivation of LET-23 EGFR in the AC is a mispositioning of the AC at the vulval midline. During vulval induction in L2 larvae, polarized LIN-3 secretion from the AC acts as an attractive cue for the VPCs (Grimbert et al., 2016; Mereu et al., 2020), with the closest VPC (P6.p) migrating underneath the AC and adopting the 1° cell fate. At the same time, P6.p produces the neuropeptide-like ligand NLP-26 which creates a positive feedback by further polarizing LIN-3 secretion towards P6.p (Mereu et al., 2020). Our results suggest that after the initial positioning of the AC relative to the VPCs has been achieved during cell fate specification, LIN-3 secreted by the 1° descendants of P6.p stabilizes the AC at the vulval midline by activating LET-23 EGFR signaling in the AC (**Fig. 7**). LIN-3 expression has been detected in the vulF cells from the L4.0 stage onward (Saffer et al., 2011), and a recent study found expression of EGL-38, the transcription factor inducing *lin-3* expression in the 1° vulval cells, as early as at the Pn.px stage in mid-L3 larvae (Rajakumar and Chamberlin, 2007; Webb Chasser et al., 2019). Together, these findings suggest that a reciprocal LIN-3 signal from the 1° vulval cells, originally identified by Chang et al. (1999), may act already before the onset of vulval lumen formation as a guidance cue for the AC. This hypothesis is consistent with the slight delay in BM breaching and enhancement of the *unc-6(lf)* AC invasion defect after inactivation of LET-23 in the AC (Ziel et al., 2008). Sherwood et al. (2003) have previously postulated the existence of one or multiple additional invasion cues from the 1° vulval cells, possibly sent as diffusible signals acting at a distance (Sherwood and Sternberg, 2003). We thus propose that LIN-3 may be one of several, redundant guidance factors expressed by the 1° vulval cells. If the AC is not correctly localized at the vulval midline prior to invasion, it cannot focus its actin-rich invadopodia at the mid-point between the 1° vulF cells, which may explain the reduced efficiency in BM breaching (Hagedorn et al., 2014). Accordingly, global inactivation of *lin-3 egf* by RNAi not only causes a vulvaless phenotype, but also resulted in a reduced number, slower assembly and longer live time of the AC invadopodia (Lohmer et al., 2016). Since the BMs were breached in around 40% of *let-23AcKO; unc-6(ev400lf)* double mutants at a later stage (L4.4), additional guidance cues besides LIN-3 EGF and UNC-6 Netrin may exist to promote BM breaching by the AC.

**Figure 7.**
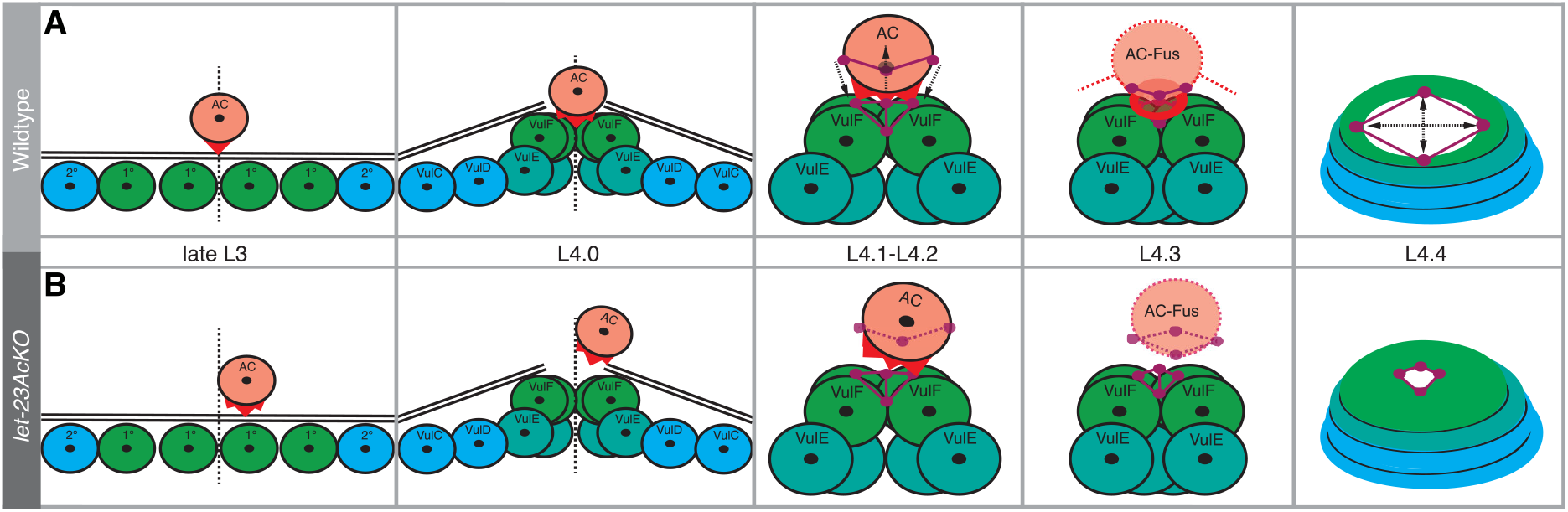
LET-23 signaling in the AC controls formation of the uterine-vulval connection. (**A**) The AC connects the ventral uterus to the vulval toroids. This process involves AC alignments and invasion at the vulval midline (L3 to L4.0) and the formation of ring-shaped adherens junctions at the AC-vulF contact site (L4.1-L4.2), allowing dorsal lumen expansion (L4.3-L4.4). (**B**) Inactivation of LET-23 EGFR in the AC perturbs AC alignment, adherens junction organization and dorsal lumen opening.

### LET-23 signaling is necessary to connect the vulva to the uterus

After breaching the BMs at the vulval midline, the AC organizes the formation of the dorsal vulval lumen (Estes and Hanna-Rose, 2009). The AC needs to physically occupy the space at the vulval midline, where the dorsal lumen will form, until it fuses with the uterine seam cell to stabilize the vulval-uterine connection (Estes and Hanna-Rose, 2009; Sapir et al., 2007).

Using microfluidic-based long-term imaging (Berger et al., 2021), we could for the first time continuously capture the whole process of vulval morphogenesis form VPC invagination until eversion within the same animal. This in-depth look into organogenesis revealed that the AC induces a complex rearrangement of newly forming adherens junctions that first constrict and then expand at the AC-vulF contact site (**Fig. 7**). Our data further suggest that LET-23 signaling in the AC organizes the formation of a circular actin ring, which may provide a scaffold for the formation of the connection between the uterus and vulva. The ring-shaped adherens junctions at the AC to vulF contact site may be necessary to subsequently expand the dorsal vulval lumen, and allow other uterine cells, such as the uv1 cells, to connect to the dorsal vulval toroids. The delayed expansion of the dorsal lumen observed after inactivation of LET-23 therefore may also be due to defects in the specification of the uv1 cells.

There are numerous studies of developmental processes demonstrating that signaling by the EGFR or other RTKs reorganizes cell junctions and the connecting actomyosin cytoskeleton. For example, activation of VEGF signaling in endothelial cells leads to the phosphorylation of VE-cadherin, which re-organizes the adherens junction and dissociates the α/β-catenin complex that normally anchors the actin cytoskeleton to adherens junctions, resulting in increased endothelial permeability and enhanced transendothelial migration of leukocytes (Gavard et al., 2008; Sidibé and Imhof, 2014). Moreover, EGFR-dependent PI3K activation after mechanical stress promotes cell contractility mediated by the formation of new focal adhesions and ROCK-dependent activation of myosin II (Muhamed et al., 2016), and EGFR signaling during ommatidia morphogenesis in *Drosophila* is necessary for myosin II-dependent apical junctional remodeling (Robertson et al., 2012). Therefore, LET-23 EGFR signaling in the AC may play a similar role, by reorganizing the adherens junctions at the AC-vulF interphase and by controlling actomyosin-mediated cell shape changes.

In summary, our data indicate that LET-23 EGFR signaling in the AC is necessary to align the developing vulva to the uterus and organize newly forming adherens junctions connecting these two organs.

## Materials and Methods

### Worm Maintenance

All strains used in this study were grown under standard conditions as described (Brenner, 1974). Animals were maintained on NGM plates seeded with OP50 *Escherichia coli* bacteria as food source at 20°C, unless noted otherwise. The N2 Bristol strain was used as wild-type. The following alleles and transgenes were used in this study:

LGI: *hmr-1(cp21[hmr-1::gfp + LoxP])* (Marston et al., 2016).
LGII: *let-23(zh131[FRT::let-23::FRT::GFP::LoxP::FLAG::let-23])* (Konietzka et al., 2020), *bqSi294[P_hsp16.41>_FRT::mCherry::his-58::FRT::GFP::his-58, unc-119(+)]* (Muñoz-Jiménez et al., 2017).
LGIII: *zhIs146[P_ACEL-pes-10>_2xNLS-FLP-D5, unc-119(+)]* (this study), *unc-119(ed3)* (Maduro and Pilgrim, 1995), *oxTi444[ttTi5605 + NeoR(+) + unc18(+)]* (Frøkjær-Jensen et al., 2014).
LGIV: *qyIs10[P_lam-1>_lam-1::GFP, unc-119(+)](Ziel* et al., 2008),
LGV: *qyIs50[P_cdh-3>_mCherry::moeABD, unc-119(+)]* (Ziel et al., 2008), *qyIs127[P_lam-1>_lam-1::mCherry, unc-119(+)]*(Ihara et al., 2011),
LGX: *unc-6(ev400)* (Hedgecock et al., 1990), Extrachromosomal array: *zhEx614[P_ACEL-pes-10>_2xNLS-FLP-D5, P_myo-2>_mCherry]* (this study).

See **suppl. Tab. S1** for genotypes of all strains used in this study.

### Device fabrication

Short-term imaging devices were fabricated following standard soft- and photolithographic protocols (Xia *et al*. 1998). Master molds were fabricated on silicon wafers (Si-Wafer 4P0/>1/525±25/SSP/TTV<10, Siegert Wafer, Germany) using SU8 photoresist (GM1050 and GM1060, Gersteltec, Switzerland). Layers of different height were fabricated on the same wafer and aligned to each other using a mask aligner (UV-KUB3, Kloe, France). Masters for the L3s devices were fabricated with heights of 5 and 20 *μ*m, and the L4s devices at heights of 5 and 22 *μ*m. All wafers were treated with chlorotrimethyl silane (Sigma Aldrich, Switzerland) prior to PDMS casting, to prevent PDMS adhesion. Devices were fabricated from a single approximately 4 mm layer of PDMS (Elastosil RT601; ratio 10:1, Wacker, Germany) and cured at 70°C for at least 1 hour. Once hardened the device was carefully removed from the wafer, cut to size, access holes punched (15G Catheter Punch, Syneo, USA) and the device bonded to a cover glass (75×25 mm cover glass with selected thickness 0.17±0.01 mm, Hecht Assistant, Germany) using air plasma (Zepto Plasma cleaner, Diener, Germany). Long-term imaging devices were produced as described by Berger *et al*. (Berger *et al*. 2021).

### Image acquisition

All images were acquired on an upright/inverted microscope (either a BX61, Olympus, Switzerland or Ti-U, Nikon, Switzerland), equipped with a fluorescence light source (either UHP-T-460-DI and UHP-T-560-DI, Prizmatix, Israel or LedHUB, Omicron Laserage Laserprodukt GmbH, Germany), a brightfield LED (either pE-100wht, Coolled, UK or MCWHLP1, Thorlabs, USA) and a camera (iXon Ultra 888, Andor Oxford Instruments, UK or Prime95B, Photometrics, USA). z-stacks were acquired using a piezo objective drive (either Nano-F100, Mad City Labs, USA or MIPOS 100 SG, Piezosystems Jena, Germany). For confocal imaging, a spinning disk (XLight V2, Crest, Italy) was attached to the Olympus microscope, with the fluorescence LEDs coupled into the system via a 5 mm LLG. Image acquisition was controlled using custom built Matlab scripts (Matlab 2019b, Mathworks, USA) and custom built microcontrollers (Arduino Mega 2560, Arduino, Italy) for coordination of fluorescence and brightfield LEDs, piezo and camera.

Images were acquired using either a 60X water immersion lens (CFI Plan Apo VC 60XC WI NA1.2, Nikon, Switzerland), a 60X oil immersion lens (UPlanAPO 60X/1.40 Oil, Olympus Switzerland or a CFI Plan Apo Lambda 60X Oil NA1.4, Nikon, Switzerland) or a 100X oil immersion lens (UPlanSAPO 100X/1.40 Oil, Olympus Switzerland). All images were acquired at 20±0.5°C, with temperature controlled by the room air conditioning system.

### Image preprocessing and deconvolution

For analysis, the images of multiple parallel acquired worms were first cropped to the vulval region using a custom Matlab script and sorted according to their developmental stages using DIC optics or fluorescent reporter signals, such as the *qyIs50[P_cdh-3>_mCherry]* AC marker (Ziel et al., 2008), as landmarks. Fluorescent images were processed with the Huygens deconvolution software where indicated (SVI, Centre for Microscopy and Image Analysis, University of Zürich).

### Determination of developmental stages

Developmental sub-stages of L4 animals (L4.0 through L4.4) were assigned using morphological features observed with DIC microscopy, as described in (Mok et al., 2015) and by assessing fluorescent reporters such as *hmr-1::gfp(cp21), qyIs50[P_cdh-3>_mCherry::moeABD] and FRT::let-23::FRT::gfp(zh131)*.

### Quantification of AC positioning and polarity

The adherens junction marker HMR-1::GFP (*cp21*) (Marston et al., 2016) was used to identify the landmarks shown in **Fig. 3C** and quantify AC to vulF alignment with a semiautomated script in Fiji (Schindelin et al., 2012). The distance between vulA1 and vulA2, the vulF mid-point and AC borders indicated by asterisks were used to calculate the relative alignment ratio R_A_ and the absolute AC to mid-vulF distance Δ in *μ*m, as shown in **Fig. 3C**. R_A_ was calculated by dividing the distance from the AC midpoint to the nearest vulA junction (either vulA1 or vulA2) by the distance between vulA1 and vulA2. Dorso-ventral AC polarity was measured in summed z-projections of the in *qyIs50[P_cdh-3>_mcherry::moeABD]* reporter signal, as described by Mereu et al. (2020).

### Scoring AC invasion

BM breaching was scored in mid-L3 (Pn.px, Pn.pxx), early (L4.0) and mid-L4 (L4.4) larvae, as indicated in **Fig. 4**. The continuity of the BM was evaluated by fluorescence microscopy using the *qyIs127[P_lam-1>_lam-1::mCherry]* transgene.

### Quantification of AC dynamics

L4.0 stage animals were loaded into the short-term imaging device (see **Extended methods** section), and z-stacks (spacing 0.25 *μ*m) of the *qyIs50[P_cdh-3>_mcherry::moeABD]* reporter outlining the AC were acquired every 30 seconds for a total of 10 minutes. Up to three animals were imaged in parallel at 60X magnification. Acquired images were first deconvolved, roughly cropped and registered using the StackReg and TurobReg Fiji plugins (Thévenaz et al., 1998), followed by maximum intensity projection. In a second step images were centered on the brightest dot found in the AC, further cropped to a 144×90 pixel image and thresholded, yielding a binary image outlining the AC. Using the Fiji-based Image CorrelationJ plugin (Chinga and Syverud, 2007) the correlation indices for consecutive time points in a series were calculated to assess the degree, by which AC shape changes over time. The correlation index C_I_, used as a measure for AC dynamics, was calculated as the average of all correlation indices in a time series.

### Long-term imaging vulval morphogenesis

Animals carrying the *hmr-1::gfp(cp21)* and *qyIs50[P_cdh-3>_mcherry::moeABD]* reporters were trapped in the long-term imaging devices (type L2-A, Berger et al. 2021) and z-stacks (spacing 0.25 *μ*m) were acquired every fifteen minutes for a total of 48 hours from the early L3 stage until the L4.9 sub-stage (vulval eversion). After cropping, z-stacks of each timepoint were registered using the StackReg and TurobReg Fiji plugins (Thévenaz et al., 1998) to correct for possible motion during image acquisition, deconvolved and manually cropped around the vulval region. Each registered and cropped time point was 3D-projected (x-, y-, z-projections) and the time points were concatenated to generate **Movies S3** and **S4**.

### Imaging the actin cytoskeleton in the AC

Spinning disk confocal images of the *qyIs50[P_cdh-3>_mcherry::moeABD]* actin marker at the L4.3 sub-stage were deconvolved, cropped and resliced along the xz-plane to visualize the ring-shaped actin cytoskeleton at the AC-vulF interphase. 25 out of 31 control, and 18 out of 45 *let-23AcKO* animals showed a clearly defined continuous actin ring (**Fig. S5AW1**) or a semicircle (**Fig. S5AW3**), while the remaining control and *let-23AcKO* animals showed barely distinguishable and disorganized rings (**Fig. S5BW1**) or no actin ring at all (**Fig. S5BW3**). The fraction of animals containing a complete actin ring or a semi-circle is shown in **Fig. 6C**.

### Plasmid construction and single copy genome insertion

For plasmid pSS23(P*_ACEL-pes-10_::2xNLS-FLP-D5::let-858 3’UTR* in pCFJ151), the anchor cell-specific enhancer element of *lin-3* (P_ACEL_) (Hwang, 2004) coupled with the minimal *Δpes-10* promoter element was amplified from pMW87 with the primer pair OEH185/OEH187, the 1.9 kb 2XNLS-FLP-D5::let-858 3’UTR fragment was amplified from pMLS262 (Schwartz and Jorgensen, 2016) with the primer pair OEH188/OJE111. These two fragments were subcloned and subsequently the P*_ACEL-pes-10_::2xNLS-FLP-D5::let-858 3’UTR* fragment was amplified with primer pair OSS200/OSS202 and integrated into pCJF151, after digestion with AvrII/SpeI (Frøkjær-Jensen et al., 2014), by Gibson assembly cloning (Gibson et al., 2009). pSS23 was used to generate a single-copy insertion in *oxTi444[ttTi5605 + NeoR(+) + unc18(+)]; unc-119(ed3)* on LGIII using the MosSci protocol (Frøkjær-Jensen et al., 2014).

**Suppl. Table S2** contains a list of the primers used in this study.

### Statistical analysis

Statistical analysis of continuous measurements (R_A_, Δ, I_DV_, C_I_) was performed using Student’s t-test. For quantifications of discrete values (BM breaching and actin ring formation) a non-parametric Mann-Whitney test was used, as indicated in the figure legends. Prism 9.0.0 (Gaphpad) software was used for data analysis and plotting.

## Supporting information

CAD mask files

## Acknowledgements

We thank the members of the Hajnal laboratory, Leilani Miller, Peter Askjaer and Darren Gilmour for critical discussion and comments on the manuscript. We are also grateful to Erik Jorgensen for providing plasmids (Addgene plasmid #73718; http://n2t/addgene:73718; RRID:Addgene_73718) and to the *C. elegans* Genetic Center (CGC) for providing strains (Funded by NIH Office of Research Infrastructure Programs (P40 OD010440)). This work was supported by the Kanton of Zürich and grants from the Swiss National Science Foundation no.184792 and the Swiss Cancer league no. 4377-02-2018 to AH, as well as funding by ETH Zürich to AdM.

## Competing interests

The authors declare no competing or financial interests

## Author contributions

S.S., S.B., L.M. performed experiments. S.B. developed the microfluidic methods. S.S. and A.H. analyzed the data. S.S., S.B. and A.H wrote the manuscript.

## Notes

### Competing Interest Statement

The authors have declared no competing interest.

