## supplementary figures, tables and methods for "Reciprocal EGFR signaling in the Anchor Cell ensures precise inter-organ connection during *C. elegans* vulval morphogenesis"

**Supplementary information**

**Table of contents:**

1) Supplementary Figures S1-S5

2) Supplementary Movies S1-S6

3) Mask files

4) Extended methods with detailed step-by-step protocols

5) Supplementary Tables S1-S2

### Supplementary figures

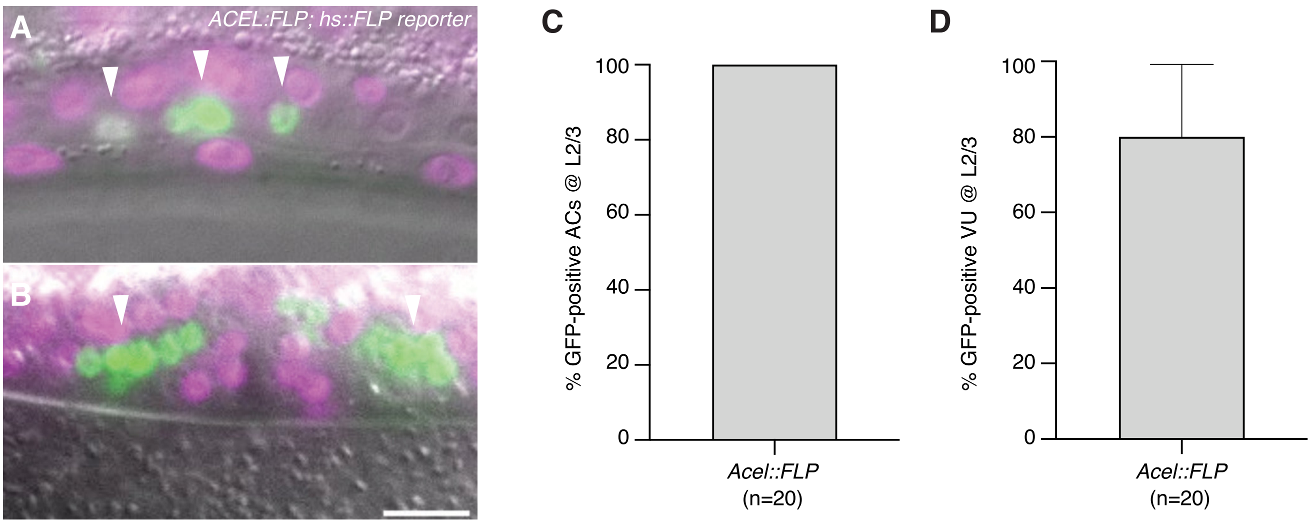


**Figure S1. Validation of the ACEL-FLP driver**

(**A-B**) Merged GFP (green) and mCherry (magenta) expression after heat-shock induction in *bqSi294[P<sub></sub>hsp16.41>FRT::mCherry::his-58::FRT::GFP::his-58]; zhIs146[PACEL-pes-10>2xNLS-FLP-D5]* animals at (**A**) the Pn.p and (**B**) the L4.4 stage. Magenta nuclei are cells without FLP activity, while FLP activity results in conversion of the *bqSi294* reporter, generating cells with green nuclei. The arrowheads in (**A**) point to the converted AC (middle) flanked by two VU cells that were also GFP positive and in (**B**) to VU descendants. (**C**) Frequency of FLP-induced conversion of the AC and (**D**) the adjacent VU cells at the L2/L3 stage. Error bars indicate the 95% CI and numbers in bracket the animals scored per condition. Scale bar is 10 $\mu$m.

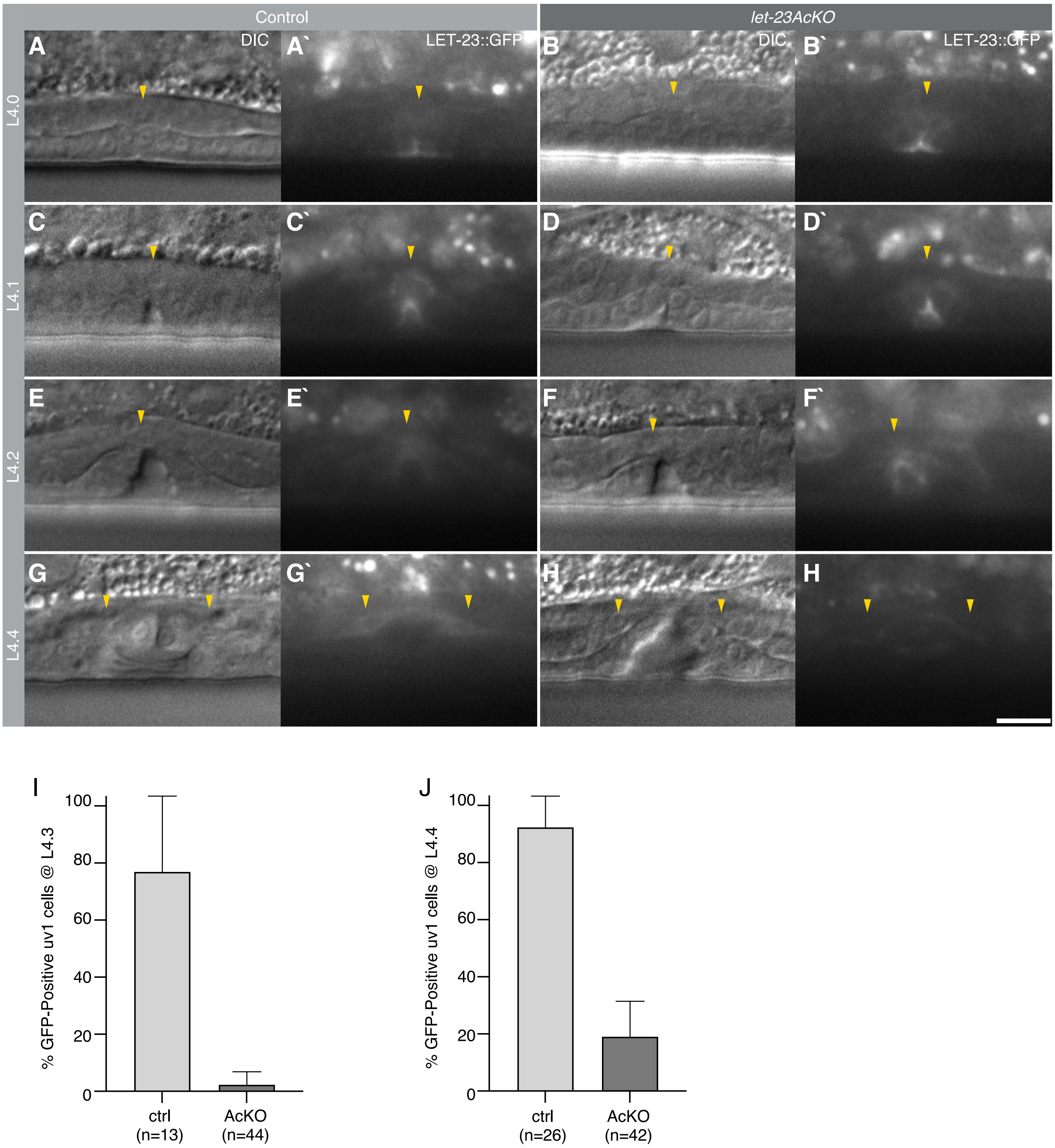


**Figure S2. Expression pattern of LET-23::GFP during vulval morphogenesis without and with ACEL-FLP-induced inactivation.**

(**A**) Control *let-23::gfp(zh131)* reporter expression at the L4.0, (**C**) L4.1, (**E**) L4.2 and (**G**) L4.4 sub-stages of vulval development. (**B,D,F,H**) FLP recombinases-mediated inactivation of *let-23::gfp(zh131)* using the *zhIs146[P*$_{ACEL\text{-}pes\text{-}10}$*>2xNLS-FLP-D5]* transgene, referred to as *let-23AcKO,* at the indicated sub-stages of vulval development. Left panels (**A-H**) show DIC images and right panels (**A'-H'**) the LET-

23::GFP signal of the mid-sagittal planes. Yellow arrowheads in (**A-F**) point to the AC and in (**G** and **H**) the uv1 cells. (**I** and **J**) Fraction of animals expressing LET-23::GFP in the uv1 cells without (ctrl) and with (AcKO) the *zhIs146[P$_{ACEL-pes-10}$>2xNLS-FLP-D5]* transgene at the L4.3 and L4.4 sub-stage. Error bars indicate the 95% CI and the numbers in brackets the animals scored per condition. Scale bar is 10 $\mu$m.

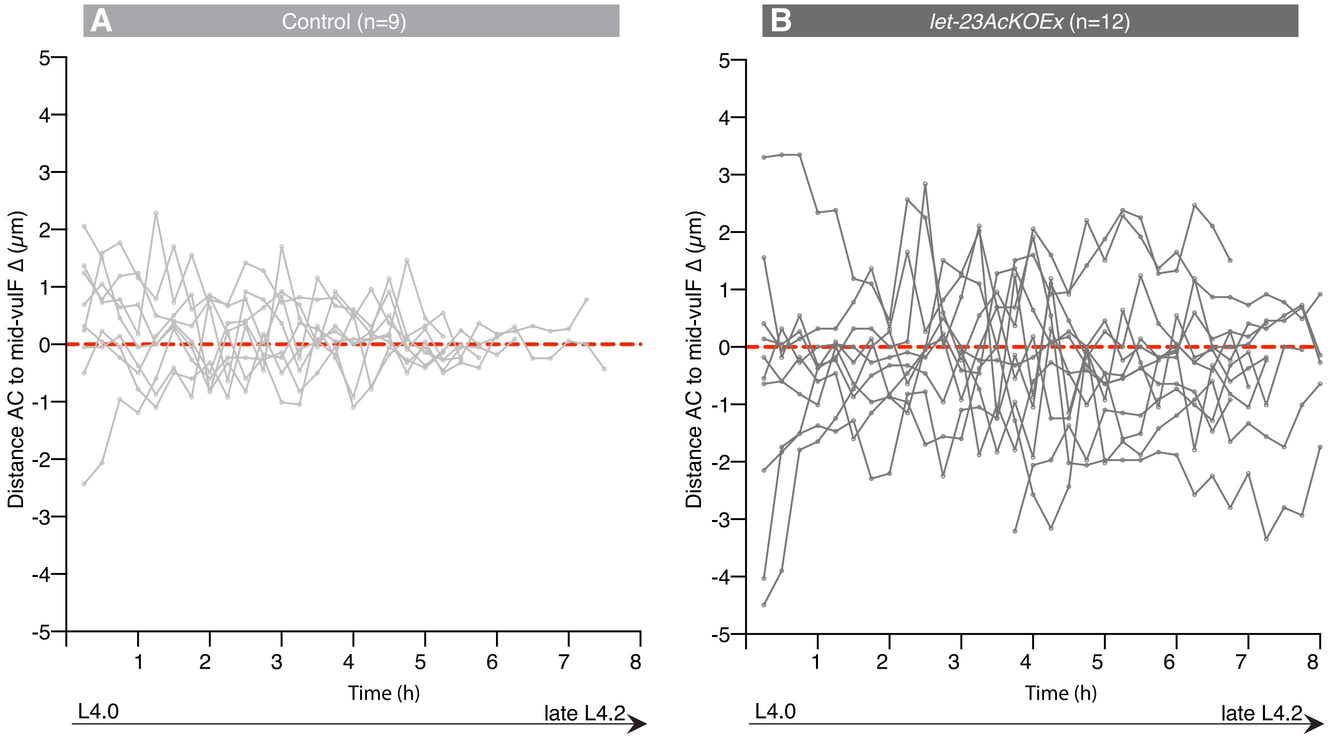


**Figure S3. AC positioning during vulval invagination**

(**A-B**)Individual measurements of the absolute AC to mid-vulF distance (Δ) in (**A**) control and (**B**) *let-23AcKOEx* animals plotted against time, used to generate the dot plots shown in **Fig. 5E,F**.

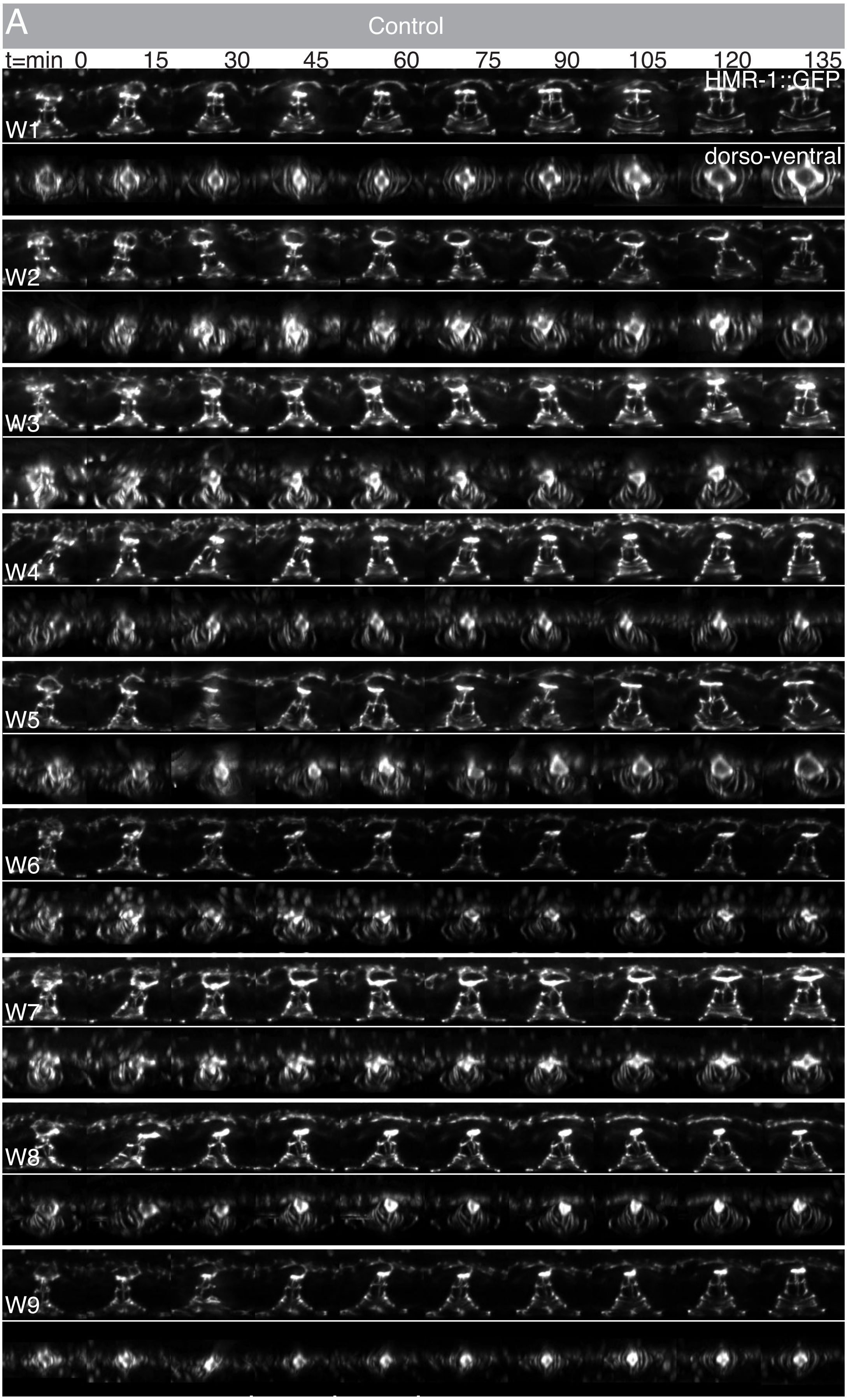
A
Control
t=min 0 15 30 45 60 75 90 105 120 135
HMR-1::GFP
dorso-ventral
W1
W2
W3
W4
W5
W6
W7
W8
W9

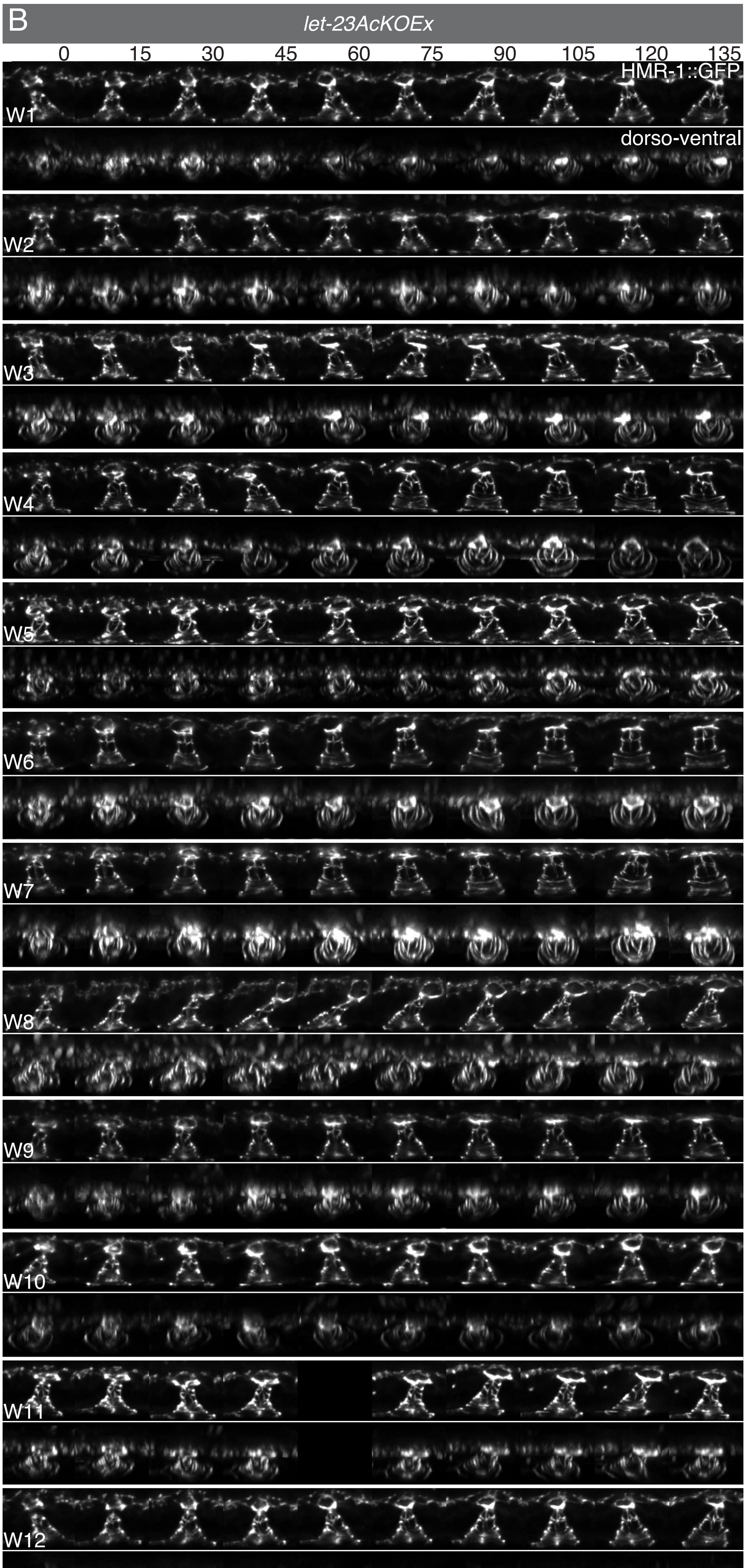
B
let-23AcKOEx
0
15
30
45
60
75
90
105
120
135
HMR-1::GFP
dorso-ventral
W1
W2
W3
W4
W5
W6
W7
W8
W9
W10
W11
W12

**Figure S4. Additional examples for toroid morphogenesis and AC-vulF junction formation over time.**

(**A-B**) Toroid morphogenesis in (**A**) nine control and (**B**) twelve *let-23AcKOEx* animals recorded at 15 minute intervals, as described in the legend of **Fig. 6**. Scale bar is 10 $\mu$m.

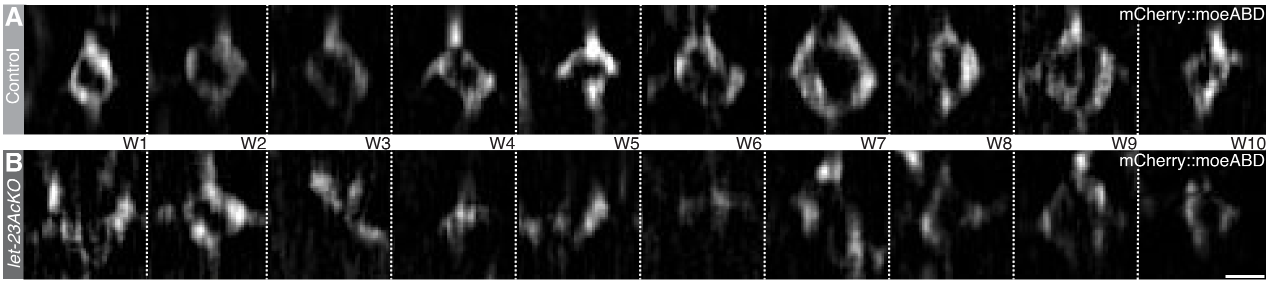


**Figure S5. Additional examples of actin ring assembly in the AC with and without LET-23 inactivation**

(**A-B**) Dorsal-ventral views (xz-projections) of the actin cytoskeleton each in ten (**A**) control and (**B**) *let-23AcKO* animals, as described in the legend of **Fig. 6D',E'**. Scale bar is 2 $\mu$m.

**Supplementary movies**

**Movie S1**: **AC cytoskeleton dynamics in a control animal.** Maximum intensity z-projections of the *qyIs50[*$P_{cdh\text{-}3}$*>mCherry::moeABD]* actin marker, acquired at 30 second intervals during the L4.0 stage.

**Movie S2**: **AC cytoskeleton dynamics in a *let-23AcKO* animal.** Maximum intensity z-projections of the *qyIs50[*$P_{cdh\text{-}3}$*>mCherry::moeABD]* actin marker, acquired at 30 second intervals during the L4.0 stage.

**Movie S3**: **Toroid morphogenesis in a control animal.** Maximum intensity z-projections of the HMR-1::GFP (*cp21*) adherens junction marker in white overlaid with the *qyIs50[*$P_{cdh\text{-}3}$*>mCherry::moeABD]* actin marker in red, acquired at 15 minute intervals from the late L3 stage on.

**Movie S4**: **Toroid morphogenesis in a *let-23AcKOEx* mutant.** Maximum intensity z-projections of the HMR-1::GFP (*cp21*) adherens junction marker in white overlaid with the *qyIs50[*$P_{cdh\text{-}3}$*>mCherry::moeABD]* actin marker in red, acquired at 15 minute intervals from the late L3 stage on.

**Movie S5**: **AC-vulF adherens junction formation and dorsal lumen expansion in a control animal.** Dorsal view (xz-projections) of the HMR-1::GFP (*cp21*) adherens junction marker at the AC-vulF interphase, acquired at 15 minute intervals between the L4.2 an L4.4 stages.

**Movie S6**: **AC-vulF adherens junction formation and dorsal lumen expansion in a *let-23AcKOEx* mutant.** Dorsal views (xz-projections) of the HMR-1::GFP (*cp21*) adherens junction marker at the AC-vulF interphase, acquired at 15 minute intervals between the L4.2 and L4.4 stages.

**Mask Files**

LOW HEIGHT.dxf CAD file for the low height structure. This structure is the same in all device types and is fabricated first. Once exposed and hardened the structures outlined by this mask are developed and the second layer is spincoated. Note the crosshairs along the midline of the mask, which serve as alignment marker for the following mask. During alignment, make sure there is sufficient overlap between the low height and higher second structure (top and bottom). Overlap on the worm side should not be too large to avoid worms slipping from one channel into the neighboring one, but also not too small to allow sufficient access to food. Aim is an overlap of 20-35 $\mu$m.

L3s.dxf CAD file for the L3s device. Eight functional units are formed on one wafer. Note the windows along the midline of the mask, which serve as alignment marker for the following mask. During alignment, the sample is viewed through the window in this mask and the crosshair formed by the previous mask is placed at the center of the windows formed here.

L4s.dxf CAD file for the L4s device. Eight functional units are formed on one wafer. Note the windows along the midline of the mask, which serve as alignment marker for the following mask. During alignment, the sample is viewed through the window in this mask and the crosshair formed by the previous mask is placed at the center of the windows formed here.

### Extended methods with detailed step-by-step protocols

#### Device Fabrication – Short term imaging.

Short-term imaging devices were made from silicon wafers (Si-Wafer 4P0/>1/525±25/SSP/TTV<10, Siegert Wafer, Germany) patterned with SU8 photoresist, following standard protocols. Briefly, in a first step wafers were cleaned with an air plasma (Diener electronic) followed by spin-coating (WS-650-23B, Laurell) the low height layer, separating inlet from outlet area, using a low viscosity SU8 (GM1050, Gersteltec). The height of this layer was chosen to be between 5 µm, high enough for liquid to pass but low enough for worms to be blocked from entering. This layer was baked and patterned using a first photomask (high-resolution film mask, Microlitho), followed by post-baking and development. In a second step, the higher SU8 layer (20 µm) was created (GM1060, Gersteltec) baked and exposed through a second photomask. Features created by the first and second mask were aligned using a mask aligner (UV KUB3, Kloe). Following exposure, wafers were post-baked and developed resulting in the final master mold. Prior to use the wafer was baked at 200°C for 10 minutes, resulting in a smooth surface and higher long-term stability of the features. The hard-baked master mold was then treated with chlorotrimethyl silane vapor (Sigma) to passivate the surface.

For device fabrication an approximately 5 mm thick layer of PDMS (PDMS Elastosil RT601 A/B, ratio 10:1) was cast on the master mold. The PDMS was then degassed under vacuum and placed in an oven at 70°C. After approximately 1 hour, the PDMS was removed from the oven and carefully peeled of the master mold. The PDMS was then cut to size access hole punched for each device (G15 puncher, Syneo) and finally bonded to cover glass (#92101400660, Glaswarenfabrik Karl Hecht) using an air plasma (Zepto, Diener electronic).

#### Step-by-step protocol

1. Clean wafers using air plasma.
2. Spincoat first SU8 layer, followed by soft-bake at 95°C.
3. Expose first layer, followed by post-bake at 95°C and development.
4. Repeat plasma cleaning after wafer is developed and dry.

5. Spincoat second SU8 layer followed by soft bake at 95°C (if necessary first bake at 65°C).
6. Expose second layer after carefully aligning the second mask to the existing features using alignment markers on either side of the wafer.
7. Post-bake and develop the wafer, followed by hard-bake at 200°C.
8. Treat wafer with chlorotrimethyl silane for at least 2 hours (NOTE: The silanes is toxic and releases corrosive vapors. This step therefore needs to be performed in a fume hood).
9. Prepare a batch of 40 g PDMS pre-polymer (Elastosil RT601 part A) and thoroughly mix it with the 4 g of crosslinker (ratio 10:1, Elastosil RT601 part B). Ideally mixing is performed using a planetary centrifugal mixer (FlackTek Speedmixer) or if not available by hand using a plastic spatula or glass rod.
10. Place the wafer in an aluminium dish and cast the PDMS onto it. Degas the wafer for about 10 minutes and after degassing break all remaining bubbles using a pipette.
11. Bake device for 2 hours at 70°C.
12. Remove the PDMS and cut it to size using a scalpel or razor blade.
13. Punch the access holes.
14. Bond the PDMS to a cover glass using air plasma and place bonded device in the oven at 70°C for a few hours.

**Worm Preparation**

Worms were bleached from mixed plates, with embryos left to hatch overnight in buffer (M9 or S-Basal). Once hatched worms were filtered through a 10 µm cell strainer (#43-10010-50, pluriSelect) to remove all unhatched eggs, as well as debris from the plate or leftover corpses. The L1 larvae were centrifuged at 1300 rcf (in 15 mL Falcone tubes) to remove supernatant and washed with 5 mL of clean buffer. Centrifugation was then repeated, supernatant removed and worms transferred to a plate seeded with OP50. Once the worms reached the desired age, they were washed off the plate using M9 buffer and left to sediment. The supernatant was then removed and worms washed three times using fresh buffer. Once washed worms were left to sediment, with as much of the supernatant as possible being removed and a small amount of buffer containing 1wt% of Pluronic F127 (#P2443-250G, Sigma Aldrich) added.

**Step-by-step protocol**

1. Starting from plates with many gravid animals.
2. Wash animals of plate using M9/S-Basal buffer.
3. Add 5% NaClO and 5M NaOH to the worm suspension (200 uL/100 uL for every 1 mL of worm suspension).
4. Gently shake bleaching mix until animals begin to break apart.
5. Centrifuge bleaching mix at 1300 rcf for 1 min.
6. Remove supernatant and add an equal amount of fresh buffer.
7. Again centrifuge at 1300 rcf for 1 min.
8. Remove the supernatant and transfer the pelleted worms to a 15 mL tube with 5 mL of fresh buffer.
9. Shake overnight at 20°C.
10. Filter the worm suspension through a 10 µm cell strainer.
11. Centrifuge worm suspension at 1300 rcf for 1 min.
12. Remove supernatant and resuspend worms in an equal amount of buffer.
13. Centrifuge worm mixture again at 1300 rcf for 1 min.
14. Remove supernatant and transfer worms to NGM plates.
15. Once worms reach desired age, wash them of the plate using fresh M9/S-Basal.
16. Leave worms to sediment by gravity or pellet by centrifugation at 750 rcf.
17. Remove supernatant and add an equal amount of fresh buffer.
18. Repeat sedimentation of centrifugation step.
19. Remove supernatant and add an equal amount of fresh buffer.
20. Repeat sedimentation of centrifugation step for a final time.
21. Remove most of the supernatant.
22. Add a small amount of M9/SBasal containing 1wt% Pluronic F127, for a final concentration of 0.1-0.5%. (NOTE: Pluronic F127 is nontoxic and only serves as lubricant during worm loading, the precise concentration is therefore is not crucial.)

23. Leave worms in the tube until worm loading.

**Worm loading**

Refer to Figure 2. Begin by preparing a 15-20 cm long piece of 1/16” tubing (#0642002, Tygon tubing 1/16’’, Fisher Scientific) connected to a 1 mL syringe, filled with M9/SBasal buffer containing 1% Pluronic F127, via a blunt needle (#300-35-970, Distrelec). Gently suck worms from the tube into the 1/16” tubing, care must be taken not to aspirate worms into the syringe. Once a sufficient amount of worms is gathered, the tubing is directly stuck into the device inlet and worms are loaded into the device channels by gently applying pressure to the syringe plunger, filling as many channels as possible. Care must be taken not to apply excessive pressure, pushing worms past the low height region into the outlet section of the device. Once loaded the tubing may be gently removed and the device mounted on the microscope. If necessary, immobilization may be improved by anesthetizing worms on chip by flushing in a solution of 100 mM Tetramisol hydrochloride (#SLBB6382V, SIGMA) in M9/SBasal buffer. For this fill a 1 mL syringe with the tetramisol solution and connect a short piece of 1/16” tubing to it, using a blunt needle. Fill the tubing with liquid, and attach it to the worm inlet. Gently apply pressure until the liquid on-chip is exchanged for the anesthetic, then remove the tubing. If an inverted microscope is used the device may be directly mounted. If an upright microscope is used it may be necessary to place the device on a carrier glass slide with the PDMS facing the carrier slide, and the cover glass facing the objective. For optimal illumination in DIC images a long working distance condenser may be necessary.

**Step-by-step protocol**

1. Start by filling a syringe with M9/SBasal buffer containing 1wt% Pluronic F127.
2. Attach a 23G needle and a long piece of 1/16” tubing to the syringe.
3. Fill the tubing with buffer from the syringe.
4. Aspirate worms from the tube into the tubing. Note do not pull the worms into the syringe and make sure no air bubbles remain in the tubing/syringe. If necessary, resuspend worms in the tube by flushing in a small amount of buffer from the syringe or tapping the tube.

5. Push the tubing into the device inlet.
6. Begin loading by gently applying pressure to the syringe buffer and monitoring device loading on a dissection scope.
7. Once loaded remove the tubing.

Optional Following Worm Loading:

8. Fill a syringe with M9/SBasal buffer containing 100 mM Tetramisol hydrochloride.
9. Attach a short piece of 1/16” tubing to the syringe using a 23G blunt needle.
10. Fill the tubing with liquid.
11. Attach the tubing to the worm inlet.
12. Gently apply pressure on the syringe to exchange the liquid on chip for the anesthetic.
13. Remove the tubing.

### Supplementary Table S1: List of strains used.

| Strain | Genotype |
|---|---|
| AH5059 | *let-23(zh131[FRT::let-23::FRT::GFP::LoxP::FLAG::let-23])II* |
| AH5158 | *let-23(zh131[FRT::let-23::FRT::GFP::LoxP::FLAG::let-23])II; qyIs50[Pcdh-3>mCherry::moeABD, unc-119(+)]V; zhEx614[PACEL-pes-10>2xNLS-FLP-D5, Pmyo-2>mCherry]* |
| AH5786 | *hmr-1(cp21[hmr-1::gfp + LoxP])I; let-23(zh131[FRT::let-23::FRT::GFP::LoxP::FLAG::let-23])II; qyIs50[Pcdh-3>mCherry::moeABD, unc-119(+)]V; zhEx614[PACEL-pes-10>2xNLS-FLP-D5, Pmyo-2>mCherry]* |
| AH5957 | *bqSi294[Phsp16.41>FRT::mCherry::his-58::FRT::GFP::his-58, unc-119(+)]II; zhIs146[PACEL-pes-10>2xNLS-FLP-D5, unc-119(+)]III; [unc-119(ed3)]III* |
| AH5958 | *let-23(zh131[FRT::let-23::FRT::GFP::LoxP::FLAG::let-23])II; zhIs146[PACEL-pes-10>2xNLS-FLP-D5, unc-119(+)]III; [unc-119(ed3)]III* |
| AH5960 | *hmr-1(cp21[hmr-1::gfp + LoxP])I; let-23(zh131[FRT::let-23::FRT::GFP::LoxP::FLAG::let-23])II; zhIs146[P$_{ACEL-pes-10}$>2xNLS-FLP-D5, unc-119(+)]III; [unc-119(ed3)]III; qyIs50[Pcdh-3>mCherry::moeABD, unc-119(+)]V* |
| AH5964 | *let-23(zh131[FRT::let-23::FRT::GFP::LoxP::FLAG::let-23])II; zhIs146[PACEL-pes-10>2xNLS-FLP-D5, unc-119(+)]III; [unc-119(ed3)]III; qyIs127[Plam-1>lam-1::mCherry, unc-119(+)]V* |
| AH5967 | *let-23(zh131[FRT::let-23::FRT::GFP::LoxP::FLAG::let-23])II; qyIs127[Plam-1>lam-1::mCherry, unc-119(+)]V* |
| AH5989 | *let-23(zh131[FRT::let-23::FRT::GFP::LoxP::FLAG::let-23])II; qyIs127[Plam-1>lam-1::mCherry, unc-119(+)]V; unc-6(ev400)X* |
| AH5990 | *let-23(zh131[FRT::let-23::FRT::GFP::LoxP::FLAG::let-23])II; zhIs146[PACEL-pes-10>2xNLS-FLP-D5, unc-119(+)]III; [unc-119(ed3)]III; qyIs127[Plam-1>lam-1::mCherry, unc-119(+)]V; unc-6(ev400)X* |
| AH6008 | *hmr-1(cp21[hmr-1::gfp + LoxP])I; let-23(zh131[FRT::let-23::FRT::GFP::LoxP::FLAG::let-23])II; qyIs50[Pcdh-3>mCherry::moeABD, unc-119(+)]V* |
| NK361 | *qyIs23[cdh-3>mCherry::PH; unc-119(+)]II; qyIs10[lam-1::gfp; unc-119 (+)]IV* (Ziel et al., 2008) |

### Supplementary Table S2: List of oligonucleotides used.

| Primer | Sequence (5`-3`) |
|---|---|
| OEH185 | CACCTGTGTATTTTATGCTGGTTTTTTC |
| OEH187 | GCTTTTTTGTACAAACTTGAATCAATGCCTGAAAGTTAAAAATTACAG |
| OEH 188 | CAAGTTTGTACAAAAAAGCAGGCTAAAAAATGCC |
| OJE111 | ATGAACCTTATTAAGGAAGATATG |
| OSS200 | CTTATAATACGACTCACTAGCACCTGTGTATTTTATGCTGG |
| OSS202 | GGGTACCAGAGCTCACCTAGTTGCCAAGCGAGGACAATTCTCATCG |
